# “Synergistic anticancer effect of combined use of (plumbagin, cis-Isoshinanolone, 3’-O-β-glucopyranosyl plumbagic acid) isolated from *Plumbago zeylanica*, induces cell death through apoptosis in human HepG2 cancer cells”

**DOI:** 10.1101/2025.03.21.644655

**Authors:** Mohana Thiruchenduran, Navin Alukkathara Vijayan, Sathyanarayana N. Gummadi, Gandarvakottai Senthilkumar Arumugam, Sivasitamparam Niranjali Devaraj

**Affiliations:** Department of Biochemistry, Meenakshi Ammal Dental College and Hospital, Meenakshi Academay of Higher Education and Research (Deemed to be University), Chennai 600095, India; Biogenix Life science Pvt Ltd, Hydrabad, Telangana 500005, India; SSS International Drug Discovery & Development Research Private Limited, Innovation & Entrepreneurship Sudha & Shankar innovation hub, IIT Madras, Chennai-600036, India; Applied and Industrial Microbiology Lab, Department of Biotechnology, Indian Institute of Technology Madras, Chennai 600 036, India

**Author notes:** **Corresponding Authors** Sivasithaparam Niranjali Devaraj, Department of Biochemistry, University of Madras, Guindy Campus, Chennai, TN 600025, India., Gandarvakottai Senthilkumar Arumugam, International Phytomedicine Innovation Private Limited, IIT Madras Research Park, Kanagam, Tharamani, Chennai, Tamil Nadu 600113.

**Keywords:** *Plumbago zeylanica*, *Cis*-Isoshinonolone, pancreatic cancer cells, cytotoxic activity, HepG2 cells

## Abstract

*Plumbago zeylanica* L., a traditional medicinal plant, has shown remarkable anti-cancer properties, making it imperative to delve into its chemical composition and biological effects. Fractions extracted from the roots of *P. zeylanica* exhibited cytotoxic activity, prompting further investigation through bioassay-guided fractionation and mass-directed isolation. This led to the discovery of three compounds: plumbagin **(1),** *cis*-Isoshinonolone **(2),** and 3’-O-β-glucopyranosyl plumbagic acid **(3).** The chemical structures of these compounds were elucidated through the analysis of 1D/2D NMR and MS data. Subsequent evaluation of compounds **1-3** for cytotoxic activity against human hepatoma cancer cells (HepG2) revealed potent effects, with all three compounds demonstrating IC_50_ values of < 0.5 µM. The combined action of these bioactive compounds in *P. zeylanic*a was found to possess effective anti-cancer properties, exhibiting superior cytotoxic activity compared to individual compounds. Specifically, compound **3** emerged as the most potent natural product in the series, displaying an IC_50_ value of 0.1 µM against the HepG2 cell line. In conclusion, compounds **1, 2**, and **3** from *P. zeylanica* showcase significant *in vitro* cytotoxic activity against HepG2 cells, underscoring their potential as promising candidates for further anti-cancer research and development.

## INTRODUCTION

In recent decades, there has been notable advancement in the healthcare system. Despite this progress, cancer remains a pressing global public health concern and continues to be one of the leading causes of death worldwide^1^.The rising incidence of cancer is particularly evident in Africa, Asia, and Central and South America, regions that account for more than 70% of all cancer-related deaths globally^2^.

Malignant diseases present significant challenges in their treatment through traditional therapeutic methods, primarily due to financial constraints and severe side effects associated with these treatments. Research studies suggest that phytochemicals may have a significant impact on both chemoprevention and chemotherapy^3^. Natural plant compounds have shown the ability to induce apoptosis and halt the cell cycle at G0/G1 and G2/M phases^4^. These findings highlight the potential of phytochemicals as a promising avenue for improving cancer treatment outcomes.

*P. zeylanica L*., commonly known as Ceylon leadwort, is a well-known herbal species with various medicinal properties. In Ayurveda, it is referred to as chitrak, with its roots known as chitramula. The name chitraka signifies its ability to cause discoloration to the skin when applied topically. This perennial plant is native to India and Sri Lanka and has been studied for its antioxidant, antimicrobial, anti-inflammatory, antidiabetic, antihyperlipidemic, antiulcer, and hepatoprotective properties. Researchers have found that *P. zeylanica* exhibits a wide range of beneficial activities, making it a valuable plant in traditional medicine.

This study aimed to investigate the novel compounds found in *P. zeylanica L*. roots for their potential anticancer properties. In order to gain a deeper understanding of the underlying mechanisms, a series of structural studies were conducted for testing purposes.

## RESULTS AND DISCUSSION

The hexane extract of *P. zeylanica* root was first separated using dichloromethane (DCM) and then further separated with ethyl acetate. The resulting residue was then fractionated and examined for its potential anticancer properties. It was discovered that the dichloromethane (DCM) fraction of *P. zeylanica* (DPz) displayed notable anticancer activity, leading to a deeper exploration of the mechanisms behind this activity. To delve into this further, a thorough analysis of DPz was carried out, involving purification through various chromatography techniques which resulted in the isolation of **1-3** compounds. The spectroscopic data of these compounds was then compared with existing literature values.

Compound (**1**) was isolated as a yellow crystalline material. The molecular formula was determined using MALDI-TOF MS to be *m/z* 213.0528, corresponding to the molecular formula C_11_H_10_NaO_3_ [M + Na + 2H]^+^. Infrared data at 3441 cm^−1^ revealed the presence of a hydroxyl group, while peaks at 1699, 1662, and 1644 cm^−1^ corresponded to a quinone carbonyl group. The UV-Vis spectrum of compound (**1**) showed absorption maxima at 415 nm and a shoulder at 268 nm, indicating a naphthoquinone derivative. The ^1^H, ^13^C NMR, and HSQC spectra revealed carbon resonances, including one methyl, five methine, and three quaternary carbons (see Table 1). The NMR analysis revealed specific signals for the compound, including a hydroxyl proton at δ_H_ 11.95 (1H, 5-OH), three aromatic protons at δ_H_ 7.61 (1H, H-8), 7.58 (1H, H-7), and 7.25 (1H, H-6), a methine proton at δ_H_ 6.79 (1H, H-3), and methyl protons at δ_H_ 2.18 (3H, 2-Me). Additionally, the ^13^C and HSQC NMR spectra showed two carbonyl signals at δ_C_ 184.8 and 190.2, a methyl carbon signal at δ_C_ 16.4, four methine carbon signals at δ_C_ 119.2, 124.1, 135.4, and 136.0, and four quaternary carbon signals at δ_C_ 115.1, 132.0, 149.6, and 161.1, indicating the presence of a trisubstituted benzene ring. Furthermore, HMBC correlations between the hydroxyl proton at δ_H_ 11.9 (5-OH) and carbons C-3, C-4a, C-8a, C-7, and C-4 suggested a connection between the carbonyl carbon and the benzene ring at C-4a. This was supported by UV-Vis absorption peaks at 268 nm and 415 nm. Correlations between the aromatic protons at δ_H_ 7.25, 7.58, and 7.61 and carbons C-1, C-4, C-4a, C-5, and C-8a further supported the presence of the benzene ring. Carbonyl group shows correlation with C-2, C-3, C-8, C-5, C-6 and Hydroxyl group attached proton shows correlation with C-3, C-4, C-7, C-4a, C-8a, overall, the NMR data strongly indicated that the compound (**1**) analysed was plumbagin (Figure 1). ^5–7^

**Figure 1.**
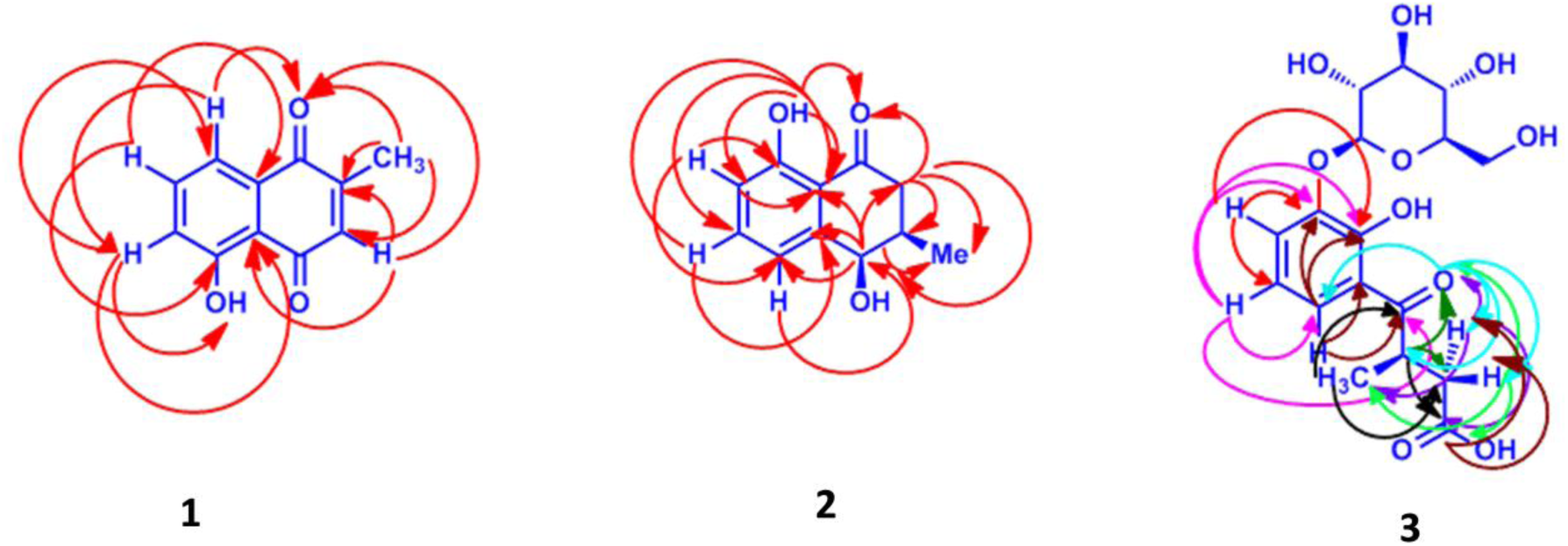
Key ^1^H−^1^H COSY and HMBC correlations of compounds 1-3.

**Table 1.** ^1^H (500 MHz) and ^13^ C (125 MHz) NMR Spectroscopic Data of Compound (1)

| Position | $\delta_{\text{C}}$ | type | $\delta_{\text{H}}$, mult. ($J$ in Hz) | COSY | HMBC |
| --- | --- | --- | --- | --- | --- |
| 1 | 184.8 | C | - |  | 2, 3, 8 |
| 2 | 149.6 | CH | - |  | - |
| 3 | 135.4 | CH | 6.79 | -<br>2-Me | 1, 4a, 2 |
| 4 | 190.2 | C | - | - | 5, 6, 8 |
| 4a | 115.1 | C | - | - | - |
| 5 | 161.1 | C | 11.95 | - | 4a, 6, 7 |
| 6 | 124.1 | CH | 7.25 | 7, 8 | 4a, 5, 8 |
| 7 | 136.0 | CH | 7.58 | 6, 8 | 5, 8a |
| 8 | 119.2 | CH | 7.61 | 5, 7 | 1, 6, 4a |
| 8a | 132.0 | C | - |  | - |
| 2-Me | 16.4 | $\text{CH}_3$ | | | 1, 2, 3 |

$^a\text{CDCl}_3$ , $\delta$ in ppm, $J$ in Hz

Compound (**2**) was isolated as a brown-colored powder. The molecular formula C_11_H_12_O_3_ was determined by HREIMS at *m/z* 233.9966, corresponding to the molecular formula C_11_H_14_NaO_4_ [M + Na + H_2_O]. The IR data at 3438 cm^−1^ indicated the presence of hydroxyl groups, with a peak at 1635 cm^−1^ corresponding to a carbonyl group. The UV spectrum of compound 2 displayed an absorption maxima at 216 nm with a shoulder peak at 258 nm, and a peak at 332 nm indicative of a tetralone derivative. The ^1^H, ^13^C NMR spectra, along with the HSQC spectrum, revealed carbon resonances attributed to one methyl, one methylene, five methine, and two quaternary carbons (Table 2). In the analysis, a downfield signal of a hydroxyl proton was observed at δ_H_ 12.3 (1H, s, H-8), along with three aromatic protons at δ_H_ 7.3 (1H, d, H-6), 6.8 (1H, d, H-5), and 6.7 (1H, d, H-5). Additionally, two methane protons were identified - one attached to a hydroxyl group at δ_H_ 4.5 (1H, brs, H-4), and another at δ_H_ 2.27 (1H, bts, 1H). Furthermore, two diastereotropic protons were detected, with one at δ_H_ 2.7 (1H, dd, H-2) and the other at δ_H_ 2.4 (1H, d, 1H). Through ^13^C and HSQC NMR analysis, a keto carbonyl at δ_C_ 205.2, a CH_2_ carbon at δ_C_ 40.5, and five CH carbons at δ_C_ 117.5 (C-7), 119.0 (C-5), 136.8 (C-6), 34.3 (C-3), and 40.5 (C-2) were identified, indicating the presence of a three-substituted benzene ring. Additionally, the proton at δ_H_ 12.5 (s, OH-8) displayed HMBC correlations to C-6, C-7, C-8a, and C-1, suggesting that the carbonyl carbon (C-1) is connected to the benzene ring at C-8a. This was further supported by UV absorptions at 332 and 258 nm. Three protons at chemical shifts of 7.3, 6.8, and 6.7 exhibit correlation with carbon shifts of 114.8, 145.3, 117.5, 119.0, and 136.8, supporting the presence of a benzene ring. The carbonyl group displays correlation with carbon shifts of 40.5 (C-2), 34.3 (C-3), and 16.01 (C-3-Me). Additionally, the hydroxyl group attached proton correlates with carbon shifts of 40.5 (C-2), 119.0 (C-5), 114.8 (8a), 145 (41), and 16.01 (3-Me). Upon careful analysis of the NMR data, it can be concluded that compound **2** is a cis-Isoshinonolone (Figure 1) ^8–9^.

**Table 2:** ^1^H (500 MHz) and ^13^ C (125 MHz) NMR Spectroscopic Data of Compound (2)

| Position | δ<sub>C</sub> | type | δ<sub>H</sub>, mult.<br>(*J* in Hz) | COSY | HMBC |
| --- | --- | --- | --- | --- | --- |
| **1** | 205.2 | C | - | - | 2,3, 3-Me |
| **2** | 40.5 | CH<sub>2</sub> | 2.72, dd, 17.0, 11.0<br>2.44, d, 17.0 | 3 | 1, 2, 3,4, 3-Me |
| 3 | 34.3 | CH | 2.27, brs | 2,2',3-Me | 2 |
| 4 | 70.6 | CH | 4.58, brs | 3,3-Me | 2, 4a, 5, 8a, 3-Me |
| 4a | 145.3 | C | - | - | - |
| 5 | 119.0 | CH | 6.83, d, 6.0 | 6,7 | 4, 8a |
| 6 | 136.8 | CH | 7.37, d, 6.0 | 5,7 | 8, 4a |
| 7 | 117.5 | CH | 6.78, d, 6.0 | 6,7 | 5, 8, 8a |
| 8 | 162.1 | CH | - |  | 1, 6, 7, 8a |
| 8a | 114.8 | C |  |  |  |
| 3-Me | 16.06 | CH<sub>3</sub> | 1.06, s |  | S, 2, 2' 3 |

<sup>a</sup>CDCl<sub>3</sub>, δ in ppm, *J* in Hz

**Table 3:**
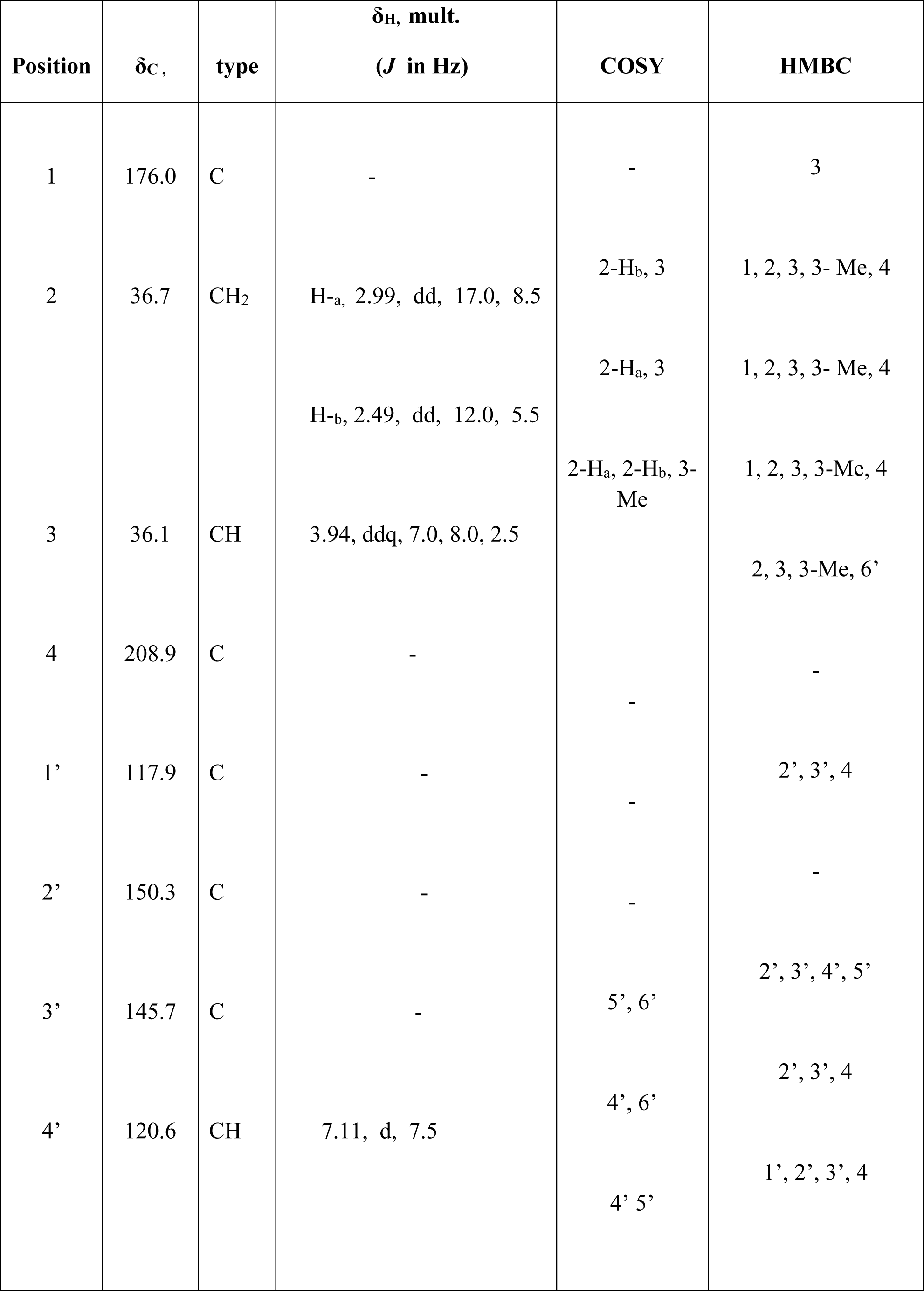
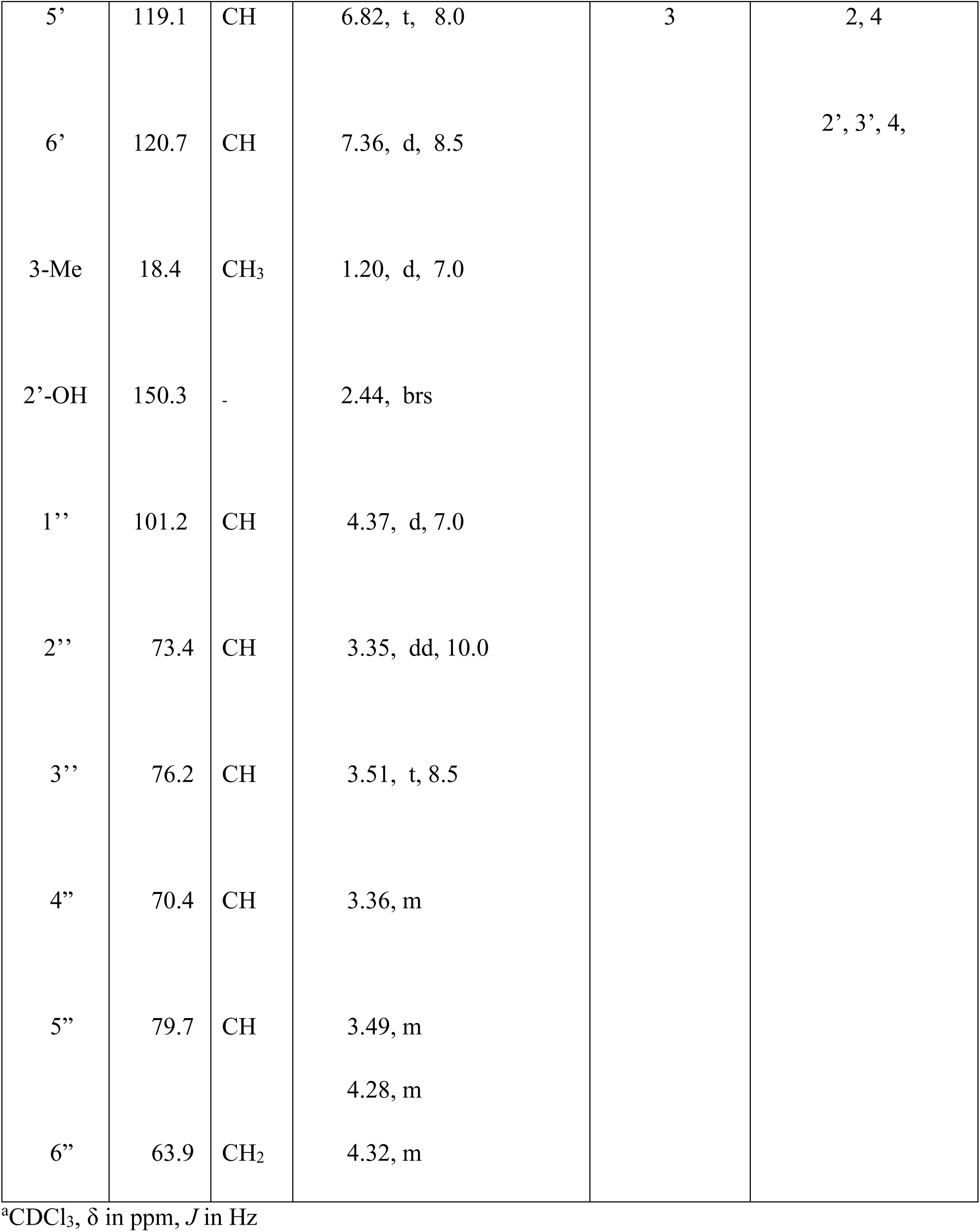
^1^H (500 MHz) and ^13^ C (125 MHz) NMR Spectroscopic Data of Compound (3)

Compound (**3**) was isolated as a brown-colored powder. The molecular formula was determined by HRESI-MS at *m/z* 391.1005, corresponding to the molecular formula C_17_H_20_NaO_9_ [M + Na-H_2_O]^+^. Additionally, a peak at *m/z* 246.9716 [M + Na-162]^+^ indicated the presence of a glycon moiety in the molecule. In the IR data, a peak at 3424 cm^−1^ revealed the presence of a hydroxyl group, while peaks at 1754 cm^−1^ (acid carbonyl), and 1637 cm^−1^ (keto carbonyl) were observed. The UV-Vis spectrum of compound (**3**) exhibited absorption maxima at 345 nm, 265 nm, and 208 nm, characteristic of a plumbagic acid derivative. Analysis using ^1^H, ^13^C NMR, and HSQC spectral techniques revealed the carbon resonances in the aglycon moiety, which included one methyl, one methylene, four methines, and three quaternary carbons. The glycon moiety was found to contain one methylene and five methine carbons. In the analysis, one can observe specific signals in the NMR spectrum of the compound. These signals include one downfield signal at δ_H_ 12.4 (1H, 2’-OH), three aromatic protons at δ_H_ 6.82 (1H, H-5’), 7.11 (1H, H-4’), 7.36 (1H, H-6’), two diastereotropic protons - one at δ_H_ 2.49 (1H, H-2b) and another at δ_H_ 2.99 (1H, H-a), one methine hydrogen at δ_H_ 3.94 (1H, H-3), and one methyl group at δ_H_ 1.2 (3H, 3-CH_3_). Additionally, a peak ranging from δ_H_ 3.35-4.37 corresponds to the glycon moiety present in the molecule, with an anomeric hydrogen at δ_H_ 4.37 indicating the attachment of a monosaccharide molecule to the ring. Further analysis through HMBC correlation reveals that the proton at δ_H_ 12.44 (2’-OH) correlates with 3’ and 4, suggesting that 4-keto carbonyls are attached to the benzene ring at C-1’. The UV-Vis absorptions at 354 nm, 265 nm, and 208 nm, along with three protons at δ_H_ 7.25, 7.58, and 7.61, show correlations with 1’, 2’, 3’, 4’, and 5’, supporting the presence of the benzene ring. The correlation of the 4-keto carbonyl with 2, 3, and 6’ and with the 3’ methyl group further indicates that the 4-keto carbonyl group is attached to the benzene ring at C-1. The presence of acid carbonyl in the compound shows a correlation with the 3-methine and 3-methyl groups. Additionally, the anomeric proton at δ_H_ 4.37 correlates with the 2’-OH group from the glycon moiety attached to the 3’-position of the benzene ring. Taking all factors into consideration, it can be concluded that compound (**3**) is 3’-O-β-glucopyranosyl plumbagic acid (Figure 1) ^10–13^.

### Cytotoxicity Studies

HepG2 cells were exposed to DPz and all three compounds at varying concentrations (1, 5, 10, 25, 50 µg/200 µl of medium) for 12, 24, 48, and 72 hours. Cell cytotoxicity was assessed using the MTT assay. The IC_50_ of DPz was determined to be 5 µg/200 µl of medium (refer to Figure 2A).

**Figure 2A:**
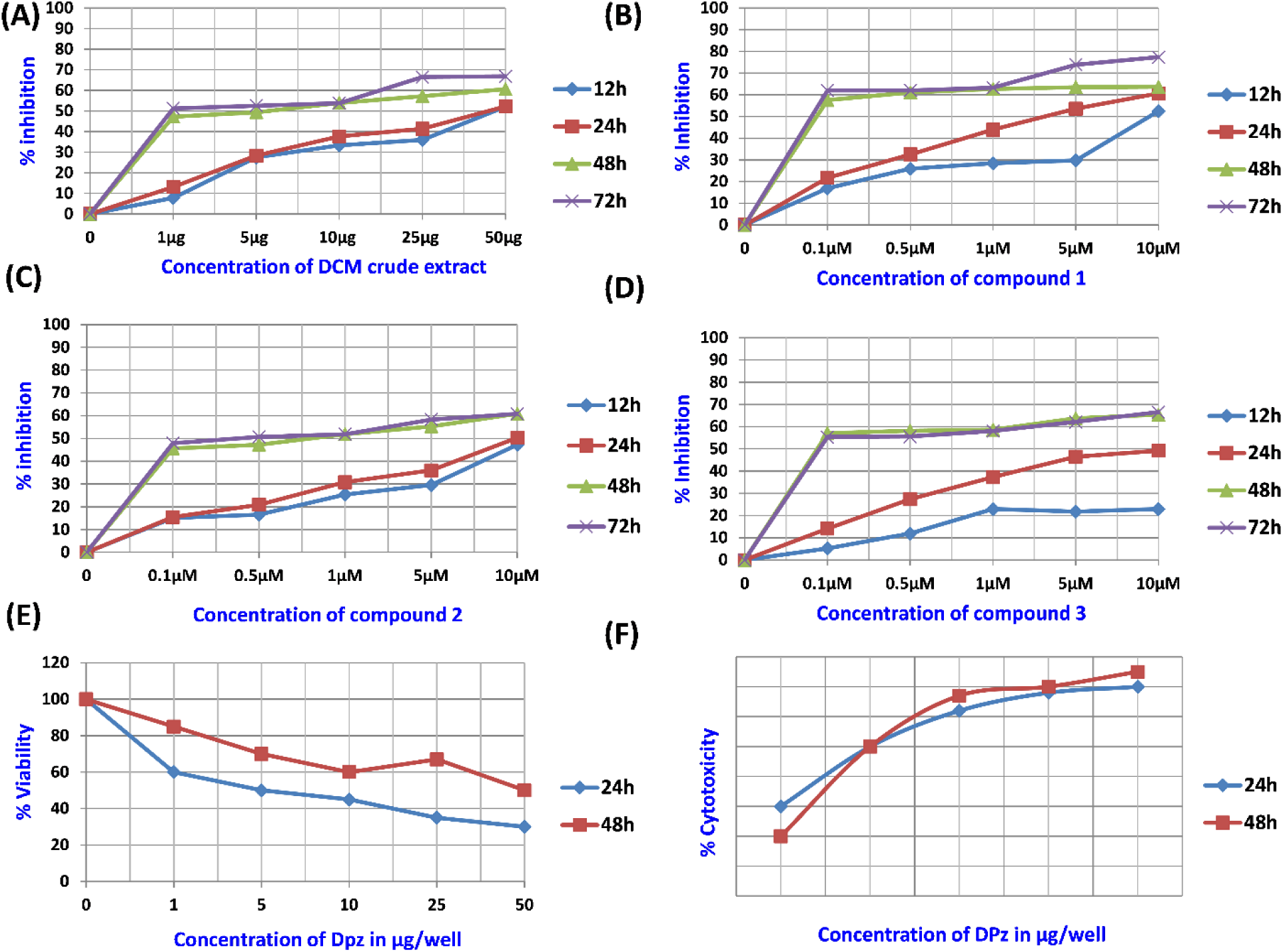
The MTT assay was conducted on DPz to assess its cytotoxic effects on HepG2 cells. The cells were exposed to DPz, which contains three compounds, at varying concentrations (1, 5, 10, 25, 50 µg/200 µl of medium) for different time intervals (12, 24, 48, and 72 hours). The cell viability was determined using the MTT assay. The IC_50_ of DPz was determined to be 5 μg/200 µl of medium. All data are presented as the mean ± standard deviation of three independent measurements. Figure 2B. The MTT assay was conducted to evaluate the effects of Compound **1** (Plumbagin) on HepG2 cells. The cells were exposed to varying concentrations of Compound A (0.1, 0.5, 1, 5, 10 µM/200 µl of medium) for different durations (12, 24, 48, and 72 hours). Cell cytotoxicity was assessed using the MTT assay. The IC_50_ of Compound **1** was determined to be 0.5μM/200µl of medium. All data are presented as the mean ± standard deviation of three independent experiments. Figure 2C. The MTT assay was conducted to evaluate the cytotoxicity of Compound **2** (Cis-isoshinonalone) on HepG2 cells. The cells were exposed to varying concentrations of Compound **2** (0.1, 0.5, 1, 5, 10 µM/200 µl of medium) for different time periods (12, 24, 48, and 72 hours). The MTT assay was used to measure cell viability. The IC50 of Compound **2** was determined to be 0.5 μM/200 µl of medium. The results are presented as the mean ± standard deviation of three independent measurements. Figure 2D. The MTT assay was conducted to evaluate the cytotoxicity of compound C (3’-O-β-glucopyranosyl plumbagic acid) on HepG2 cells. The cells were exposed to varying concentrations of Compound C (0.1, 0.5, 1, 5, 10 µM/200 µl of medium) for different durations (12, 24, 48, and 72 hours). The cell viability was assessed using the MTT assay. The IC_50_ value of compound **3** was determined to be 0.1 μM/200 µl of medium. The results are presented as the mean ± standard deviation of three independent measurements. Figure 2E. Trypan blue viability assay was conducted on HepG2 cells to assess the impact of DPz treatment at varying concentrations (1, 5, 10, 25, 50 µg/200 µl of medium) for 24 and 48 hours. The viability of cells was determined through Trypan blue viability staining. The IC_50_ of DPz was identified to be 5 μg/200 µl of medium. All data are presented as the mean ± standard deviation of three replicates. Figure 2F. LDH Leakage Assay: HepG2 cells were exposed to varying concentrations of DPz (1, 5, 10, 25, 50 µg/200 µl of medium) for 24 and 48 hours. The integrity of the cell membrane was assessed using the LDH leakage assay. The IC_50_ of DPz was determined to be 5 μg/well. All data are presented as the mean ± standard deviation of three independent measurements.

The three bioactive compounds in DPz have been discovered to possess potent anticancer properties. Compound 1, 2, and 3 exhibited IC_50_ values of 0.5 µM (Figure 2B), 0.5 µM (Figure 2C), and 0.1 µM (Figure 2D), respectively. These findings highlight the promising potential of these compounds in the field of cancer research.

Cancer is defined by the uncontrolled growth of cells and the ability to avoid programmed cell death, often by bypassing normal mechanisms of cellular apoptosis^14^. The resistance to apoptosis is a key factor in the progression of different types of cancer ^15^.

Compounds derived from plants have garnered significant attention in recent years due to their wide range of uses. Medicinal plants contain bioactive compounds that show promise in both treating and preventing various health conditions, including cancer ^16^. Research into the cytotoxic effects of phytotherapeutics for cancer treatment has sparked considerable interest, with a particular focus on the quinone-rich hexane fraction of *P. zeylanica.* This fraction has shown the most significant cytotoxicity against cancer cells, highlighting its potential as a valuable tool in the fight against cancer.

This research delves into the anticancer properties of the hexane extract of *P. zeylanica* root in human hepatic carcinoma cell lines. The results suggest that plumbagin and other compounds effectively induce apoptosis in these cells, ultimately leading to cell death through a prooxidant mechanism. Prior studies have shown that plumbagin has the potential to decrease cell viability and inhibit proliferation in different cancer types ^15–17^.

Plumbagin has been scientifically proven to be an effective anticancer agent. Researchers have found that its efficacy is primarily due to its ability to engage in redox cycling, which generates reactive oxygen species (ROS), and to chelate trace metals within biological systems.^18^ Derived from the roots of *P. zeylanica L*., plumbagin has emerged as a powerful anti-tumor compound following extensive in vitro and in vivo studies. Its effectiveness extends to the inhibition of various cancer cell lines.^19–20^

Plumbagin is primarily isolated from the Hexane extract of the root of *P. zeylanica*, yielding 95%. It is worth noting that the Hexane extract of *P. zeylanica,* containing a high concentration of Plumbagin, exhibits significantly lower anticancer activity compared to DPz extract. Studies by Qiu ^21–23^ have confirmed that Plumbagin is one of the most effective anticancer compounds derived from plant sources to date. These findings suggest that the additional compounds present in the DPz extract work synergistically with Plumbagin to enhance its pharmacological effects, making it a more effective treatment option than Plumbagin alone in the hexane extract. After considering these facts, it was decided to investigate the synergistic effect of all the isolated compounds and crude DPz. A treatment drug with an IC_50_ value of 5 µg for 48 hours was chosen for this purpose. Through our discussions and cytotoxic studies, it has become clear that the synergistic effect of all three isolated compounds present in DPz results in better cytotoxic activity compared to the individual compounds alone. This demonstrates that the anti-carcinogenic properties of DPz are not solely reliant on Plumbagin, but also on the other two compounds that work together with Plumbagin to provide the most effective defence against cancer. As a result of these findings, further studies were conducted with DPz to explore its potential benefits in cancer treatment.

### MORPHOLOGICAL STUDIES

The microscopic examination of control and treated cancer cell morphology provides direct evidence of the therapeutic potential of the drug being utilized. The morphological changes induced by DPz in cells, including rounding up, loss of cell-to-cell contact, and membrane blebbing, were clearly observed (refer to Figure 3). Cancer cells cultivated in *in vitro* cultures exhibited distinct morphological characteristics that varied among different cell lines.^24–25^

**Figure 3:**
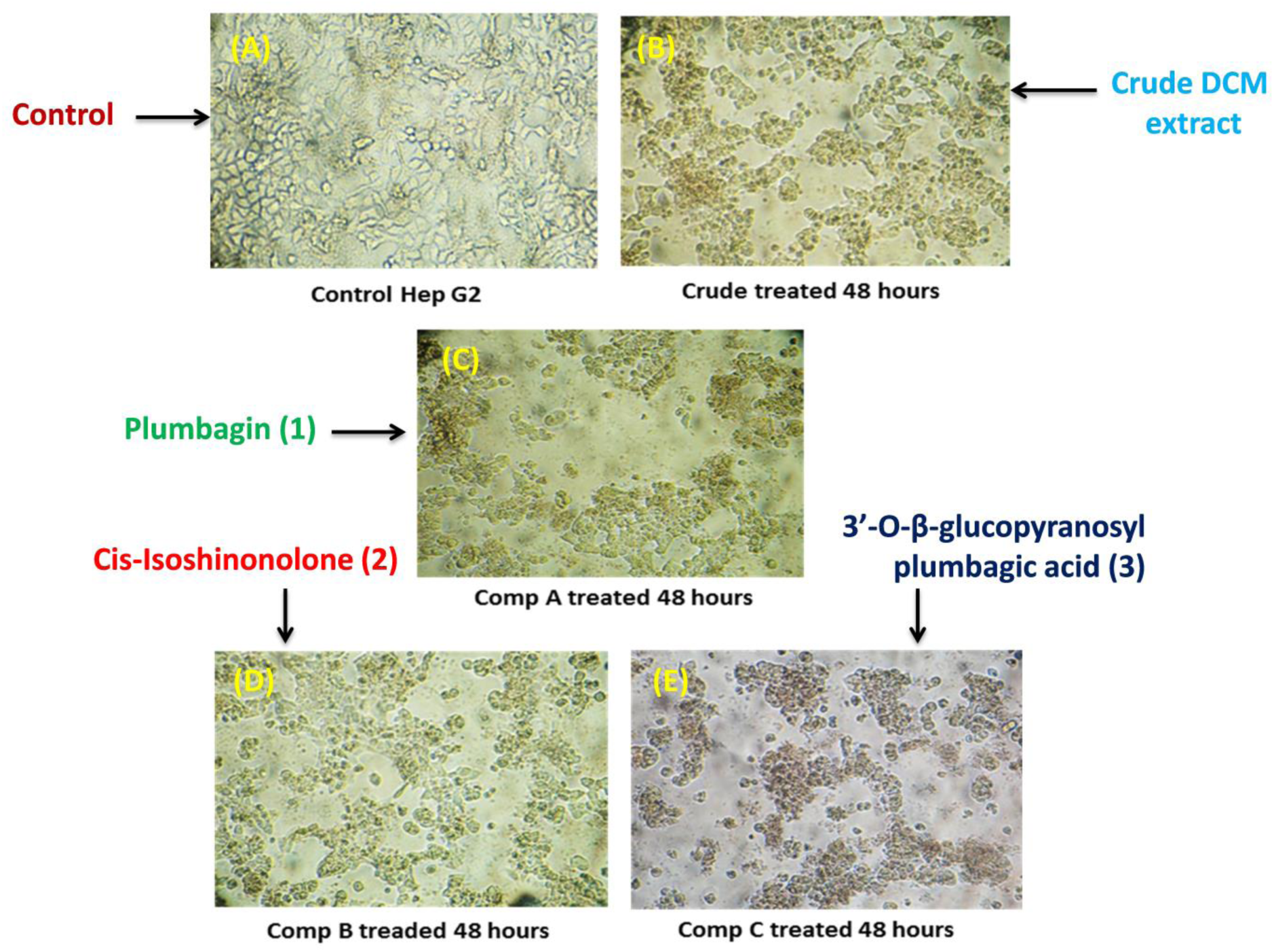
Morphological changes in control and treated HepG2 cells. It was observed that treated cells exhibited a more compact morphology compared to control cells, with a higher concentration of granules located towards the center of the cells. Following a 2-day exposure to DPz and other compounds, there was a noticeable decrease in the number of cells visible on the dish in comparison to the initial count, particularly when compared to the other compounds. Additionally, the treated cells appeared smaller, contracted, and displayed abnormal nuclear morphology.

The HepG2 cells exhibit an elongated spindle-shaped morphology. When cancer cells undergo apoptosis due to drug treatment, significant changes occur in cell morphology that can be observed under a phase contrast microscope. The breakdown of the proteinaceous cytoskeleton by caspases during apoptosis leads to cell shrinkage and rounding up.^26^ Additionally, membrane blebbing and loss of contact with adjacent cells are common features of apoptosis, as illustrated in Figure 3. An early detachment of monolayer adherent cells from their basal membrane, known as anoikis, is another characteristic of apoptosis. These cells can be observed floating in the medium^27^.

The findings of the current study have unveiled that DPz treatment induces apoptosis-related morphological changes, confirming its cytotoxic effect. The impressive anticancer potential exhibited by the DPz fraction can be attributed to the presence of bioactive compounds identified through GC–MS and NMR analysis. These compounds, such as plumbagin (**1**), cis-Isoshinonolone (**2**), and 3’-O-β-glucopyranosyl plumbagic acid (**3**), are recognized as potent phytoconstituents with significant anticancer properties.

## CONCLUSION

The potent anticancer activity DPz fraction owes to the bioactive molecules present in it. The elucidations of such compounds play a major role in understanding the pharmacological role and helps manufacturing of synthetic drugs. TLC and HPLC parameters showed that DPz fraction contained three major compounds. Silica column was used to separate and purify the compounds from DPz fraction. Each compound was subjected to Mass spectrometry and NMR studies for the elucidation of their respective structures. The first eluted Compound (**1**) was found to be Plumbagin second eluted Compound (**2**) was Cis-Isoshinonolone and third eluted compound C was 3’-O-β-glucopyranosyl plumbagic acid.

## EXPERIMENTAL SECTION

### General Experimental Procedures

Optical rotations were measured using a JASCO Polarimeter-P2000, with the [α]_D_ values reported in deg cm^2^ g^_1^. UV spectra were recorded on a Jasco V 550 UV-VIS spectrophotometer. IR spectra were obtained using a Perkin Elmer Spectrum One Fourier Transform Infrared spectrometer with KBr pellets.^1^H and ^13^C NMR spectra of compounds were recorded on a Bruker Advance 500 NMR spectrometer in CDCl_3_ with TMS as the internal standard, and chemical shifts (δ) were reported in ppm. Two-dimensional ^1^H–^1^H COSY, NOESY, ^1^H–^13^C HSQC, HMBC, and spectra were also recorded on a Bruker Avance 500 NMR spectrometer. MALDI-TOF MS analyses were conducted using an Applied Biosystems ABI4700 TOF mass spectrometer in reflector mode with an accelerating voltage of 20 kV. HRESIMS were measured on a Q-TOF micro mass spectrometer (Waters USA) in Positive ion mode using methanol as the solvent. Chromatograph HPLC (Shimadzu’s RID 10 A), Column, C_18_ 128 Shim-pack XR-ODS II (150 mm (L) x 3 mm (i.d.). Mobile Phase (A: 0.1% Acetic acid in CH_3_CN, B: 0.1% Acetic acid in H_2_O, Gradient: 2% to 10 % of CH_3_CN in 60 minutes, Flow rate: 1 ml/min, Detection: 254 nm, Injection volume: 20 µl, Column Temperature: 30 ^0^C

### Plant Material

The plant *P. zeylanica* was harvested from the hills of Thrikur village in the Thrissur district of Kerala, India in May of 2011. The specimen was authenticated by Dr. Joshy K. Simon, Head of the Department of Botany at Christ College in Irinjalakuda, Thrissur, Kerala (680125). This authentication was crucial to ensure the accurate identification of the plant species, as any error in identification could result in significant misinterpretation of the findings. The chemicals used in the study included MTT [3, (4,5-dimethylthiozal-2-yl)-2,5,-diphenyltetrazolium bromide] at a concentration of 5 mg in 1X PBS, and DMSO at a concentration of 0.1%. These chemicals were utilized in the research process to achieve reliable and reproducible results.

### Extraction and Isolation

#### Isolation of compounds from *Plumbago zeylanica* root

The Hexane extract of *P. zeylanica* root was initially fractionated using dichloromethane (DCM) and then further fractionated with ethyl acetate. The resulting residue was then fractionated and tested for its anticancer properties. It was observed that the DCM fraction of *P. zeylanica* (DPz) exhibited significant anticancer activity, prompting further investigation into the underlying reasons for this activity. To achieve this, a comprehensive analysis of DPz was conducted, involving extraction, isolation, purification, identification, and complete structural elucidation.

The dried and finely powdered roots of *P. zeylanica* (5 kg) were extracted with 5L of hexane over a period of 5 days at room temperature. The resulting hexane extract was then concentrated under reduced pressure, and the residue was further fractionated with 5L of dichloromethane over another 5-day period at room temperature. The solvent was subsequently removed under reduced pressure. The resulting dichloromethane extract (5 g) was then subjected to silica gel column chromatography using a 230-400 mesh silica gel (200 g). A gradient of ethyl acetate in n-hexane was used, and a total of 30 fractions were collected. This process allowed for the isolation and purification of various compounds present in the *P. zeylanica* roots, providing valuable insights into the chemical composition of this plant material (Figure 4-6).

**Figure 4:**
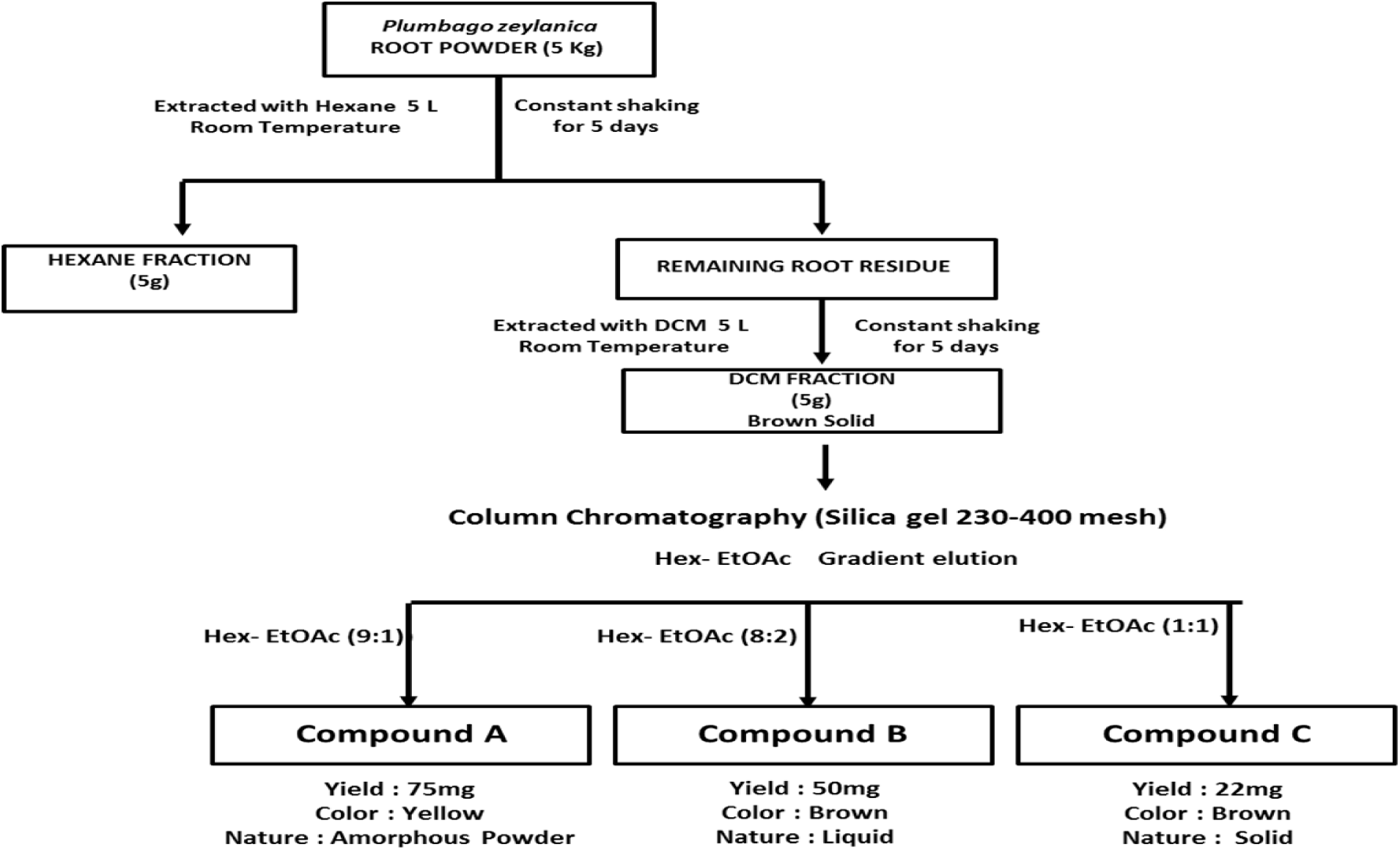
Flow chart representing the steps of compound isolation

**Figure 5.**
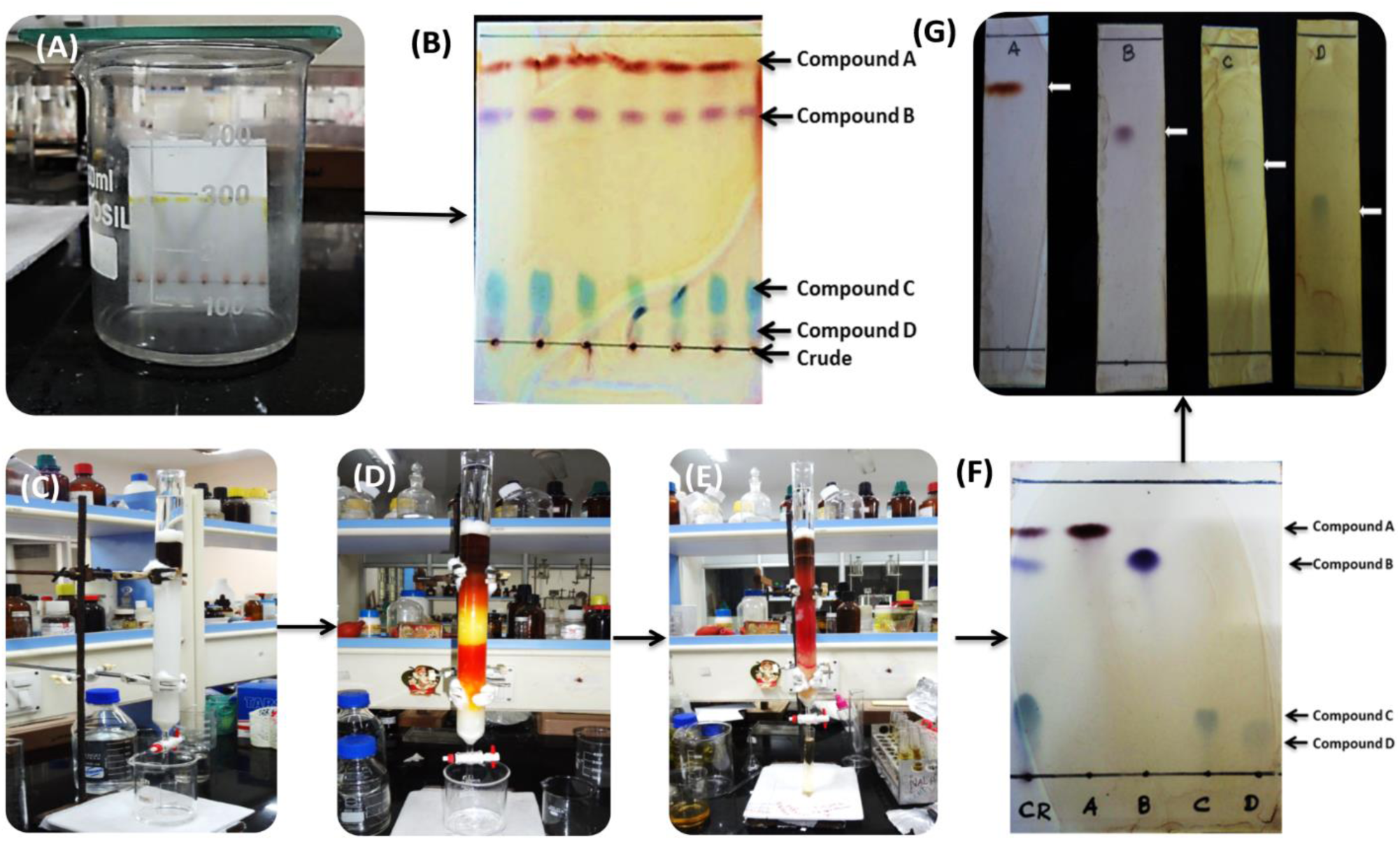
Scheme of purification of DCM fraction of *P. zeylanica* (DPz) of by column chromatography. a) TLC of crude sample (b) TLC of crude sample after run with Hexane/ Ethylacetate (60:40) solvent as eluent (D) Before elution (D) After elution (D) After complete elution (f) TLC of purified compound after run with Hexane/Ethylacetate (60:40). (G) TLC of pure single compounds using with Hexane/Ethylacetate (60:40) solvent as eluent. Column chromatography was carried out on 230-400 mesh silica gel (200 g) and Column size: (id 30m × 90 cm).The crude DCM fraction of *P. zeylanica* (DPz) was purified using column chromatography packed with a 230-400 mesh silica gel using a gradient of Hexane/Ethyl acetate (9:1 to 1:1) used as the eluent. Fractions were collected and concentrated under vacuum to afford pure compounds. The desired product was monitored in a TLC with pre coated alumina sheet silica. The isolated compounds were obtained compound (**1**) (75 mg), compound (**2**) (50 mg), compound (**3**) (22 mg).

**Figure 6A.**
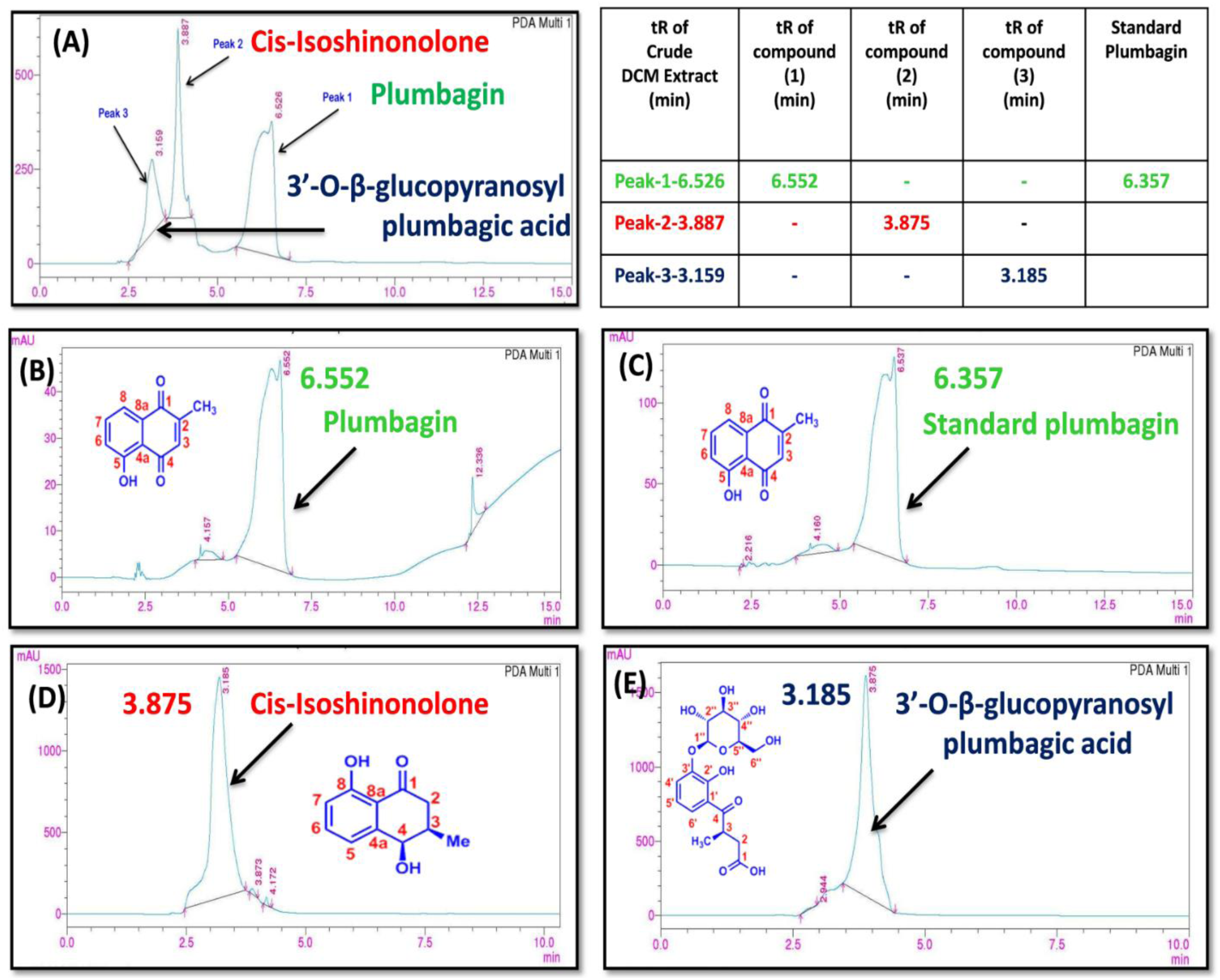
displays the HPLC chromatogram of all three compounds, while Figures B, D, and E represent the separated compounds from the DCM fraction. Figure 6C shows the standard plumbagin. Discrete HPLC analysis was conducted to assess the purity and distinct characteristics of the isolated compounds. A clear single peak was observed for all three compounds, indicating that they were pure and unadulterated. Figure 6F illustrates the retention times of the three compounds identified in the DPz. HPLC chromatogram, confirming the purity and consistency of the isolated compounds with those found in DPz. Subsequent structural elucidation was conducted utilizing various spectroscopic parameters.

Fractions (C01-10) eluted with hexane-EtOAc (9:1, v/v) afforded compound (**1**) as yellow colour needles (75 mg), mp 72 ^0^C, R_f =_ 0.8 (hexane-EtOAc, 6:4).

Fractions (C15-C20) eluted with hexane-EtOAc (8:2) afforded compound (**2**) as brown colour liquid (50 mg), Rf: 0.7 (Hexane-EtOAc, 6:4)

Fractions (C25-C30) eluted with hexane-EtOAc (7:3) afforded compound (**3**) as Brown colour powder (22 mg), Rf: 3.5 (Hexane-Ethylacetate, 60:40).

All the fractions were monitored by TLC aluminium sheet silica 60 F_254_ plates and visualisation was carried out by spraying with alcoholic Ferric chloride solution which shows coloured spots

##### Compound 1

Color: Yellow powder, Yield: 75 mg, Melting Point: 72 °C, TLC: Rf = 0.8 (Ethyl acetate-Hexane, 60:40). Solubility: This compound is soluble in Chloroform, Dichloromethane, Ethyl acetate, Methanol, Ethanol, Acetonitrile, and DMSO. UV: (MeOH) λ_max_ (log ε) 268 (0.867), 415 (0.212) nm. IR: (KBr), ν_max_ = 3441 (OH stretch), 2956 (m, CH str, sym, CH_3_), 2924 (m, CH str, asym CH_2_), 2852 (m, str, sym CH_2_), 1744, 1699, 1662, 1644 (s, Quinone Carbonyl), 1609, 1455 (s. C=C, Aromatic), 1379 (m, bending, sym, CH), 1364, 1335, 1303, 1258, 1230, 1186, 1162, 1114 (C-O str), 1084, 1051, 1021 (C-O str of OH), 983, 920 (C-H bending, Aromatic), 835, 800, 751, 671, 596, 554, 504, 464 cm^−1^.

^1^H NMR (500 MHz, CDCl_3_): δ 2.18 (3H, s, Me-2), 6.79 (1H, brs, H-3), 7.25 (1H, m, H-6), 7.58 (1H, m, H-7), 7.61 (1H, m, H-8), 11.95 (1H, OH-5).

^13^C NMR (125 MHz, CDCl_3_): δ 16.4 (2-CH_3_), 115.1 (C-4a), 119.2 (C-8), 124.1 (C-6), 132.0 (C-8a), 135.4 (C-3), 136.0 (C-7), 149.6 (C-2), 161.1 (C-5), 184.7 (C-1), 190.2 (C-4).

MALDI-TOF-MS: *m/z* (positions) 213.1081 [M+Na+2H]^+^, 191.4463 [M+Na-H_2_O]^+^,173.1830[M-CH_3_]^+^,164.5856 [M+Na-CO-H_2_O]^+^, 228.9713 [M+CH_3_CN]^+^, 251.5663 [M+CH_3_CN+Na]^+^,267.4294 [M+CH_3_CN+K]^+^, 379.9377 [2M+2H]^+^. C_11_H_10_NaO_3_ [M + Na+2H]^+^ calc. 213.0528, found. 213.1081

MALDI -TOF-MS: *m/z* (neg.ions): 188.7802 [M-H]^−^, 211.8418 [M-H+Na]^−^, 229.1712 [M-H+CH_3_CN]^−^ CHN: Anal. calcd for :C_11_H_8_O_3_, C, 70.16; H, 5.89, Found: C, 74.16; H, 5.60

##### Compound 2

Colour: Brown colour powder, Yield: 50 mg, [α]_D_^25^ : 4.5 ° (c = 0.5, CHCl_3_), TLC: R_f_ = 0.7 (Ethyl acetate-Hexane, 60:40), Solubility: Soluble in Chloroform, Dichloromethane, Ethyl acetate, Methanol, Ethanol Acetonitrile, and DMSO.

UV: (MeOH) λ max (log ε) 216 (2.252) 258 (1.214), 332 (0.546) nm,

IR: KBr), *ν*_max_ cm^−1^ : 3438 (O-H stretch), 2962 (m, CH str, sym, CH_3_) 2929 (m, CH str, sym CH_2_) 1635 (s, C=O str, Ketone), 1579, 1455 (s, C=C str, Aromatic), 1410 (s, C-C str, Aromatic), 1343, 1242, 1216 (s, C-O, Ar-O) 1165 (C-O str), 1090, 1060 (C-O of OH), 978, 941, 905,(s, C-H bend, Aromatic) 872, 813, 797, 745 (s, CH bend), 718 (CH_2_ Rocking), 621(O-H bend), 594, 479.

^1^H NMR (500 MHz, CDCl_3_): δ 1.06 (3H, brs, CH_3_), 2.27 (1H, brs, H-3), 2.44 (1H, d, *J* = 17.0 Hz, H_eq_-2), 2.72 (1H,dd *J* = 17.0, 10.5 Hz, H_ax_-2), 4.58 (1H, brs, H-4), 6.78 (1H*, J* = 6.0 Hz, H-7), 6.83(1H, *J* = 6.0 Hz, H-5), 7.37(1H, d, *J* = 6.0 Hz, H-6).

^13^C NMR (125 MHz, CDCl_3_): δ 16.0 (CH_3_), 34.3 (C-3), 40.5 (C-2), 70.6 (C-4), 114.8 (C-8a), 117.5 (C-7), 119.0 (C-5), 136.8 (C-6), 145.3 (C-4a), 162.1 (C-8), 205.2 (C-1).

HR-ESI-MS: C_11_H_14_NaO_4_ [M+Na+H_2_O]^+^ Calc. 233.2162, found. 233.9966

##### Compound 3

Colour: Brown colour powder, Yield: 22 mg, [α] ^25^: 10.4 ° (c = 0.1CHCl_3_), TLC: R_f_ = 3.5 (Ethyl acetate-Hexane, 60:40), Solubility: Soluble in Chloroform, Dichloromethane, Ethyl acetate, Methanol, Ethanol, Acetonitrile, and DMSO.

UV: (MeOH) λ _max_ (log ε) 208 (2.150) 265 (0.504), 354 (0.104) nm

IR: (KBr), *ν*_max_ = 3424 (O-H stretch), 3026, 2925 (m, CH str, asym,CH_2_) 2852 (m, CH str, sym, CH_2_), 2237, 1735 (s, C=O str, acid Carbonyl) 1637 (s, C=O, Keto Carbonly) 1492, 1452 (s, C=C str, Aromatic), 1377, 1325, 1272, 1183 (C-O-C), 1052, 1027(C-O), 912 (C-H bend Aromatic), 891, 817, 761, 701 (CH_2_ Rocking), 589,

^1^H NMR (500 MHz, CDCl_3_): δ 1.2 (3H, d, *J* = 7.0 Hz, 3-CH_3_), 2.99 (1H, dd, *J* = 17.0, 8.5 Hz, H-2a), 2.49 (1H, dd, *J* = 12.0, 5.5 Hz, H-2b), 3.94 (1H, ddq, *J* = 7.0, 8.0 Hz, 3-H), 4.37 (1H, *J* = 6.5 Hz, H-1’’), 6.82 (1H, t, *J* = 8.0 Hz, H-5’), 7.11 (1H, d, *J* = 7.5 Hz, H-4’), 7.36 (1H, d, *J* = 8.5 Hz, H-6’),

^13^C NMR (125 MHz, CDCl_3_): δ 18.5 (3-CH_3_), 36.1 (C-3), 36.7 (C-2), 63.9 (C-6’’), 70.4 (C-4’’), 73.4 (C-2’’), 76.2 (C-3’’), 79.7 (C-5’’) 101.2 (C-1’’), 117.9 (C-1’), 119.1 (C-5’), 120.6 (C-4’), 120.7 (C-6’), 145.7 (C-3’), 150.3 (C-2’), 176.0 (C-1), 208 (C-4). HRESI-MS: *m/z* (pos.ions) 391.1151 [M+Na-H_2_O]^+^, 246.9716 [M+ Na-162]^+^ C_17_H_20_NaO_9_ [M + Na-H_2_O]^+^ calc. 391.1005, found. 391.1151

### Cytotoxicity experiments

#### MTT ASSAY (Mosmann 1983)

The assay quantifies the reduction of MTT [3-(4,5-dimethylthiazol-2-yl)-2,5-diphenyltetrazolium bromide] by mitochondrial succinate dehydrogenase. MTT penetrates the cell membrane and migrates to the mitochondria, where it is transformed into an insoluble, dark purple formazan product. Subsequently, the cells are lysed using an organic solvent, and the resulting solubilized formazan reagent is quantified using spectrophotometry. This method is based on the fact that only metabolically active cells can reduce MTT, making the intensity of the formazan product a reliable indicator of cell viability.^28^

##### Procedure

Hep 3B cells were seeded in 96-well plates at a cell density of 1 × 10^6 cells per well and incubated overnight at 37°C with 5% CO_2_. Following this, the complete medium was replaced with 100 µl of incomplete medium containing various concentrations of four extracts dissolved in 0.1% DMSO. The cells were then incubated for 24 and 48 hours, respectively. After the incubation period, the medium with the drug was replaced with 100µl of incomplete medium containing 10 µl of MTT and incubated in the dark for four hours at 37°C. Subsequently, the medium with MTT was discarded, and 100 µl of DMSO was added to solubilize the dark blue formazan crystals. The plate was then placed on a rocker for 5-10 minutes. The plate was read using an ELISA plate reader (DNM-9602 microplate reader, Beijing Perlong New Technology Co., Ltd) at 570 nm, and the percentage viability was calculated accordingly.

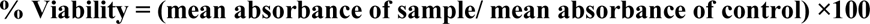

#### Trypan blue staining (Strober, 2001)

Reagents required:

Trypan blue dye (0.4% in saline)

##### Procedure

Cells treated with the extracts for various durations were isolated and diluted with medium to achieve a cell density of 2.4 x 10^6 cells per milliliter. A 10µl portion of the suspension was combined with an equal volume of 0.4% trypan blue stain and allowed to incubate at room temperature for 15 minutes. Excess stain was then removed with 1X PBS, and 10µl of the suspension was loaded onto the haemocytometer stage. After a 2-minute incubation period, the sample was examined under a light microscope. Dead cells appeared blue, while live cells remained colorless. A total of 100 cells were counted per sample, and the percentage of viable cells per sample was calculated accordingly.

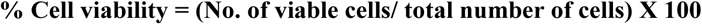

#### Assay of Lactate dehydrogenase

LDH leakage from the drug treated cells, was measured by King’s method (1965)

##### Reagents required

1. 0.1N Glycine buffer: 7.5 g of glycine and 595 mg of sodium chloride in 1 liter distilled water
2. Buffered substrate (pH-10): 2.76 g of Lithium Lactate were dissolved in 125 ml of glycine buffer containing 75 ml of 0.1N NaOH solution. This was prepared just before use
3. 0.4N NaOH solution
4. 5mg of NAD+ was dissolved in 10ml distilled water. Prepared just before use
5. DNPH 1mM: 200mg of DNPH was distilled in 1 liter of 1N HCl
6. Standard pyruvate solution: 11 mg of Sodium Pyruvate was dissolved in 100 ml of 0.1N Glycine buffer.

##### Procedure

Cells were cultured at a density of 1X10^6 and treated with varying concentrations of the drug for 12, 24, and 48 hours. Following the incubation period, the tubes were centrifuged at 3000rpm for 15 minutes to separate the supernatant and cell pellet. The cells were then lysed using a lysis buffer, and protein levels were determined using Lowry’s method.

Next, 0.1ml of the supernatant was combined with 1ml of buffered substrate and incubated at 37°C for 15 minutes. After this initial incubation, 0.2 ml of NAD+ solution was added, and the mixture was further incubated for 15 minutes. The reaction was then assessed by adding 1ml of DNPH reagent and incubating for an additional 15 minutes at 37 °C. Finally, 7 ml of 0.4 N NaOH solution was added, and the color developed was measured at 420 nm using a UV-visible spectrophotometer.

The activity of the enzyme was quantified in µ mol of pyruvate formed per minute per milligram of protein.

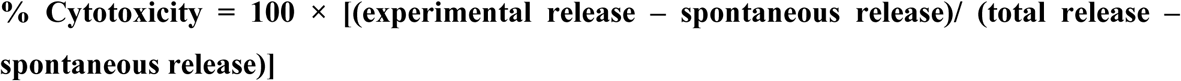

Spontaneous release – is the amount of LDH activity in the supernatant of control cells; Experimental release – is the amount of LDH activity in the supernatant of cells treated with the extracts and Total release - is the activity in cell lysate ^29^.

## STATISTICAL ANALYSIS

The data was analyzed using SPSS version 7.5 software. Statistical significance was determined using the student’s t-test, and probability values (p-values) were reported. A p-value of less than 0.001 was considered statistically significant. Results were presented as Mean ± Standard Deviation (S.D.).

## ASSOCIATED CONTENT

The Supporting Information is available free of charge at 1D and 2D NMR, MALDI-TOF and ESI-MS spectra of compounds (**1**)-(**3**)

## AUTHOR INFORMATION

### Notes

The authors declare no competing financial interest.

## ACKNOWLEDGMENTS

The authors express their gratitude to CSIR–Central Drug Research Institute, for their support. Additionally, the authors would like to extend their thanks to the SAIF, Department of Chemistry, and IIT Madras for providing access to FT-IR, NMR, and ESIMS facilities.

## Supplementary Information

### General Experimental Procedures

**Figure S1.**
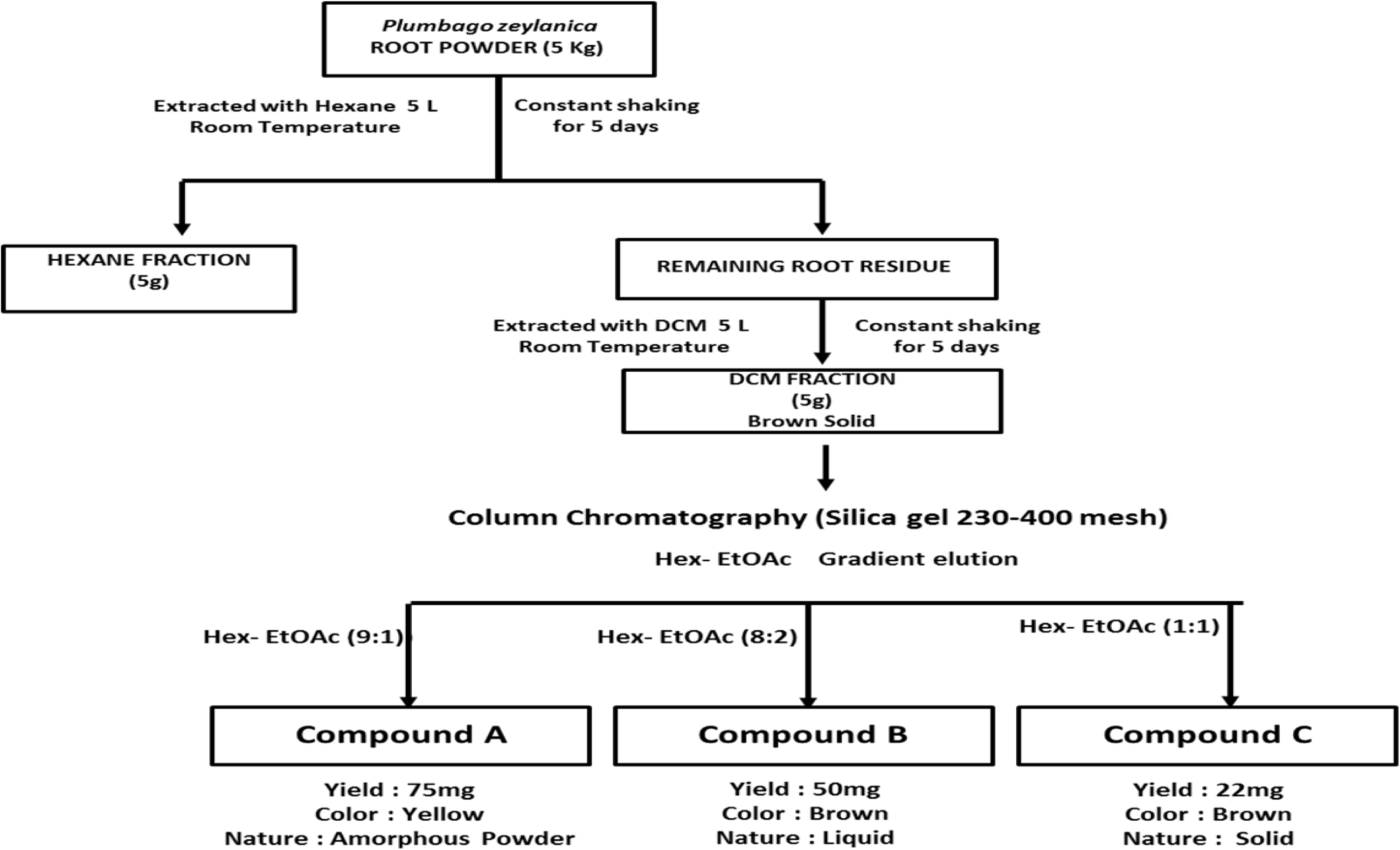
Flow chart representing the steps of compound isolation

**Figure S2.**
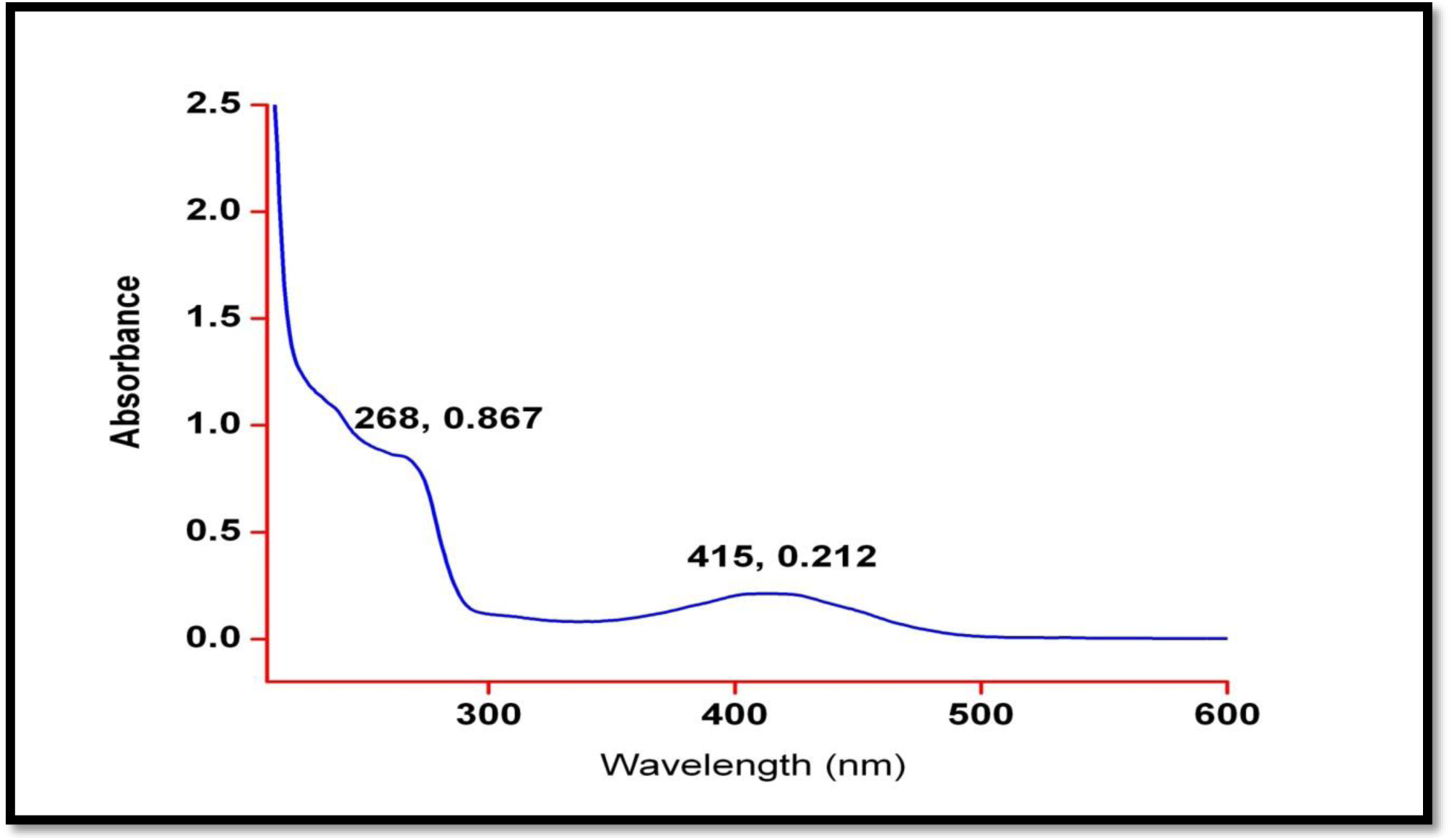
UV/Vis spectrum of plumbagin (1) in methanol. UV: (MeOH) λ max (log ε) 268 (0.867), 415 (0.212) nm

**Figure S3.**
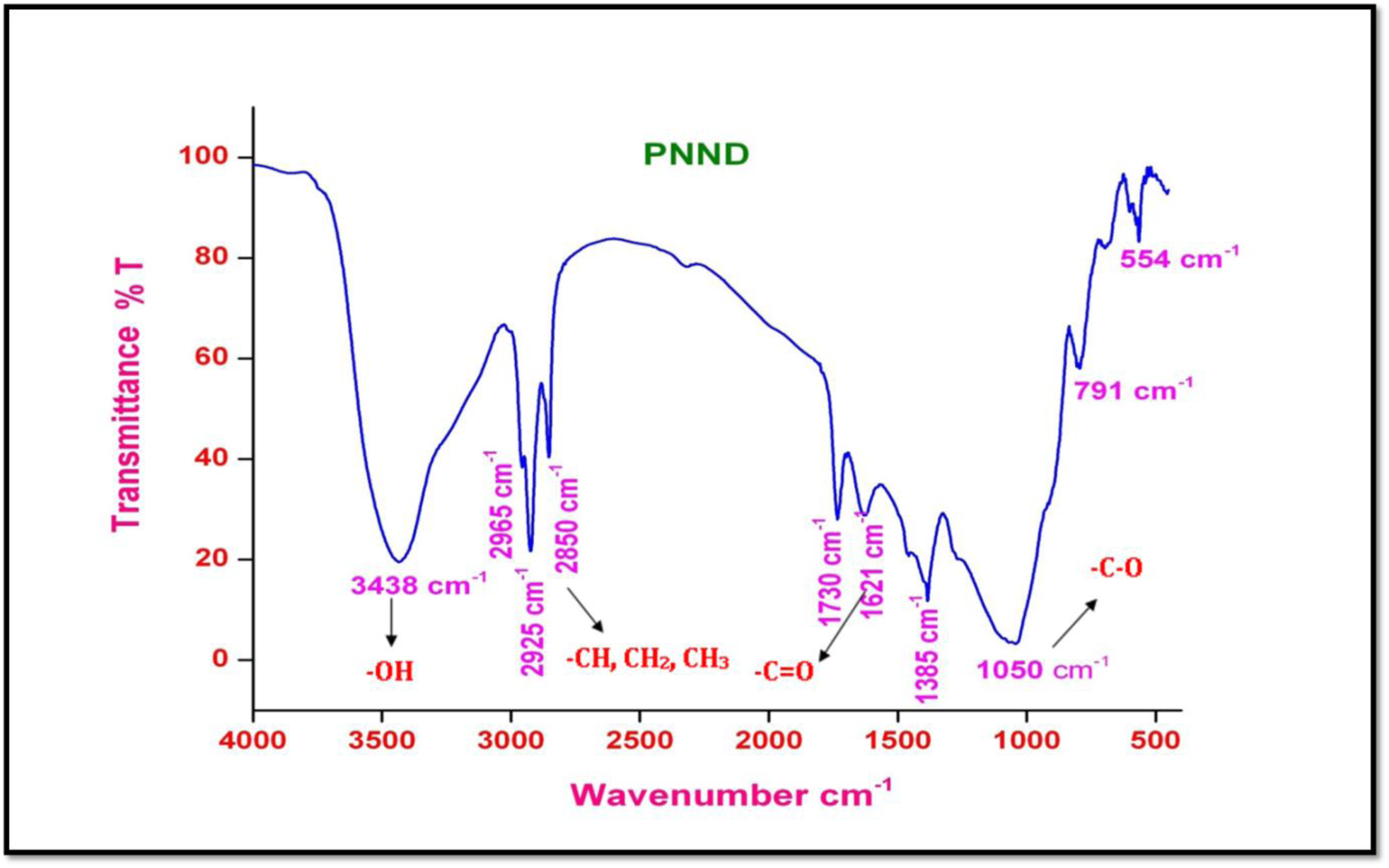
IR spectrum of plumbagin (1).

**Figure S4.**
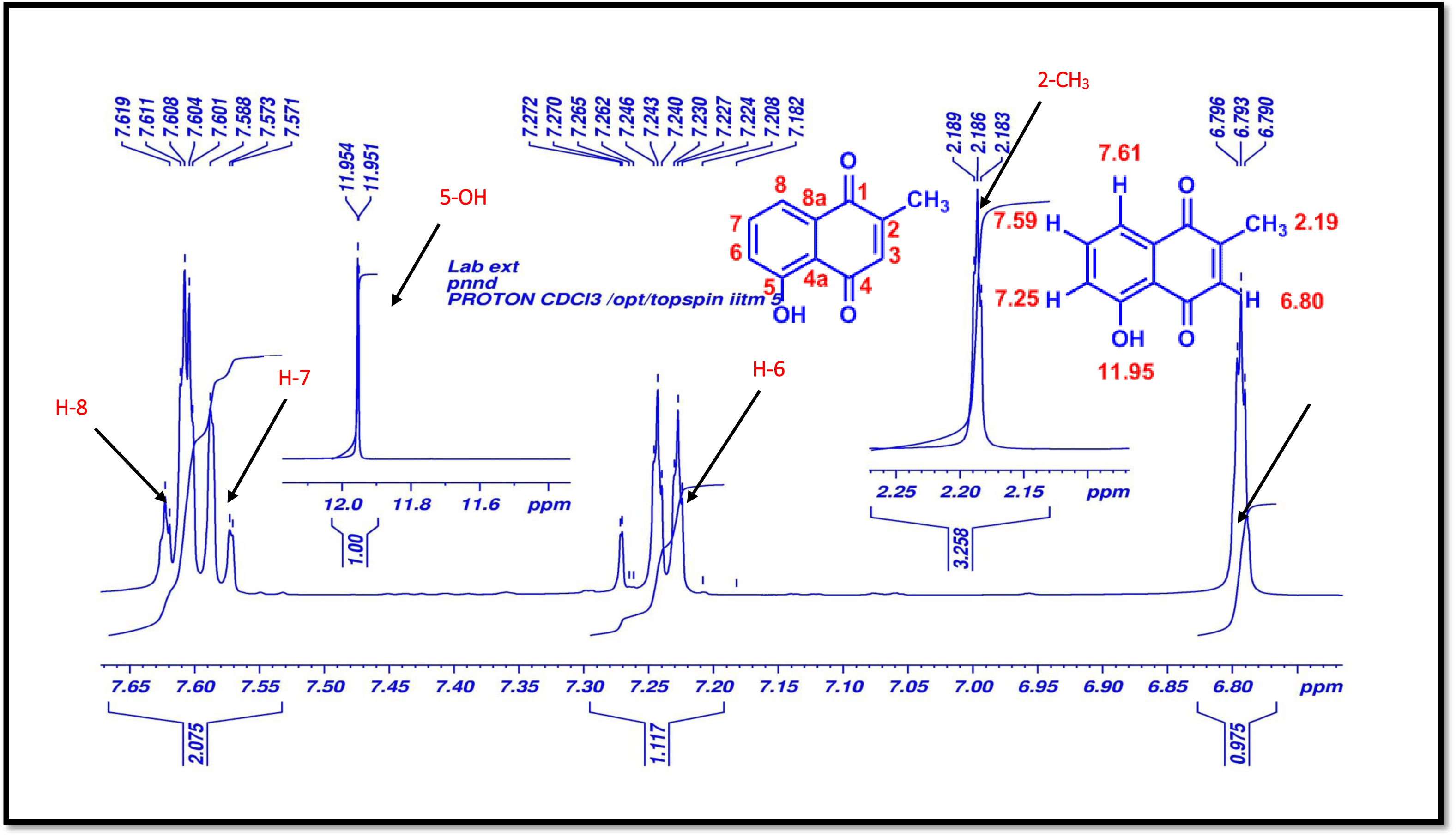
^1^H-NMR (500 MHz, CDCl_3_) spectrum of plumbagin (1).

**Figure S5.**
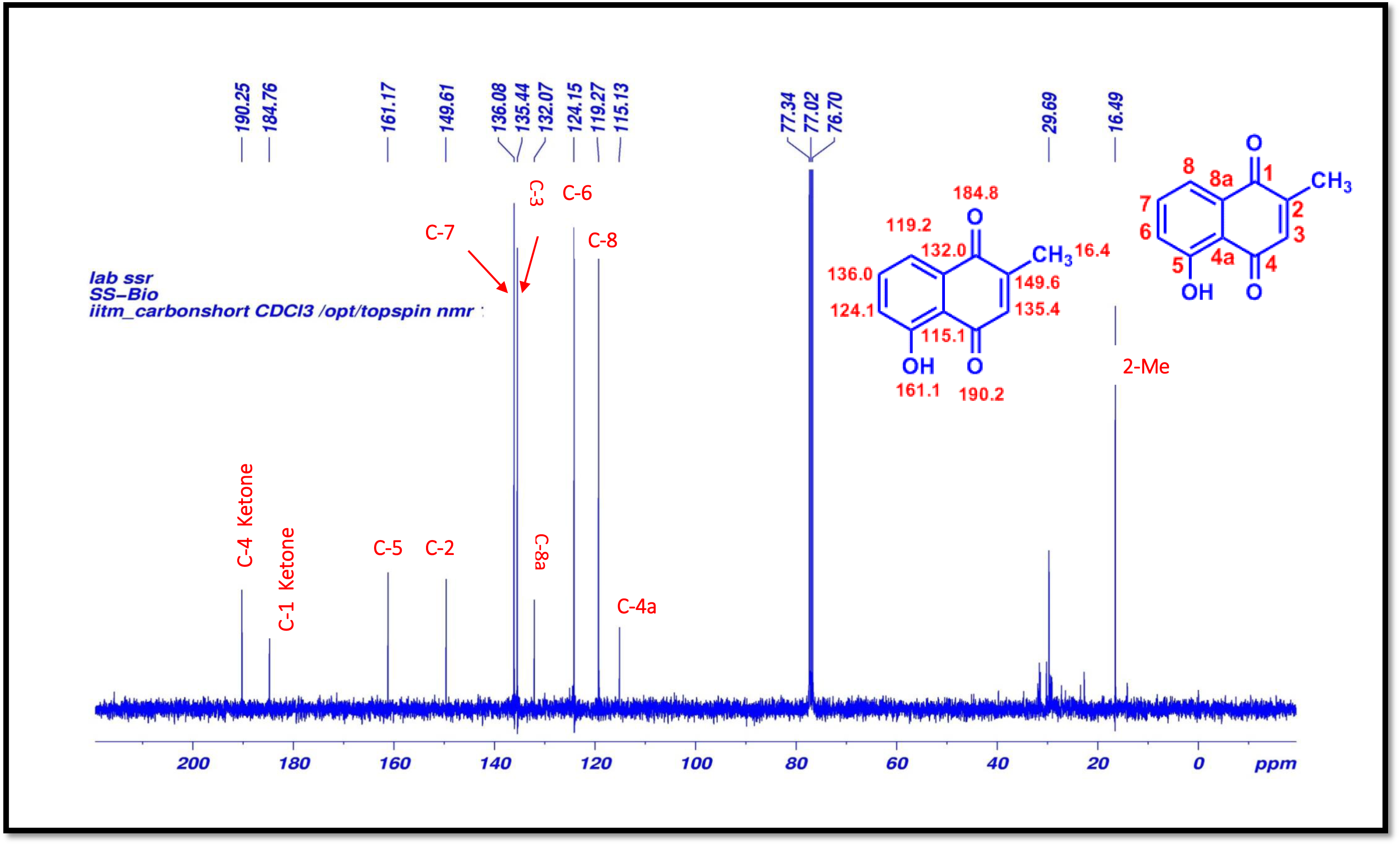
^13^C-NMR (125 MHz, CDCl_3_) spectrum of plumbagin (1).

**Figure S6.**
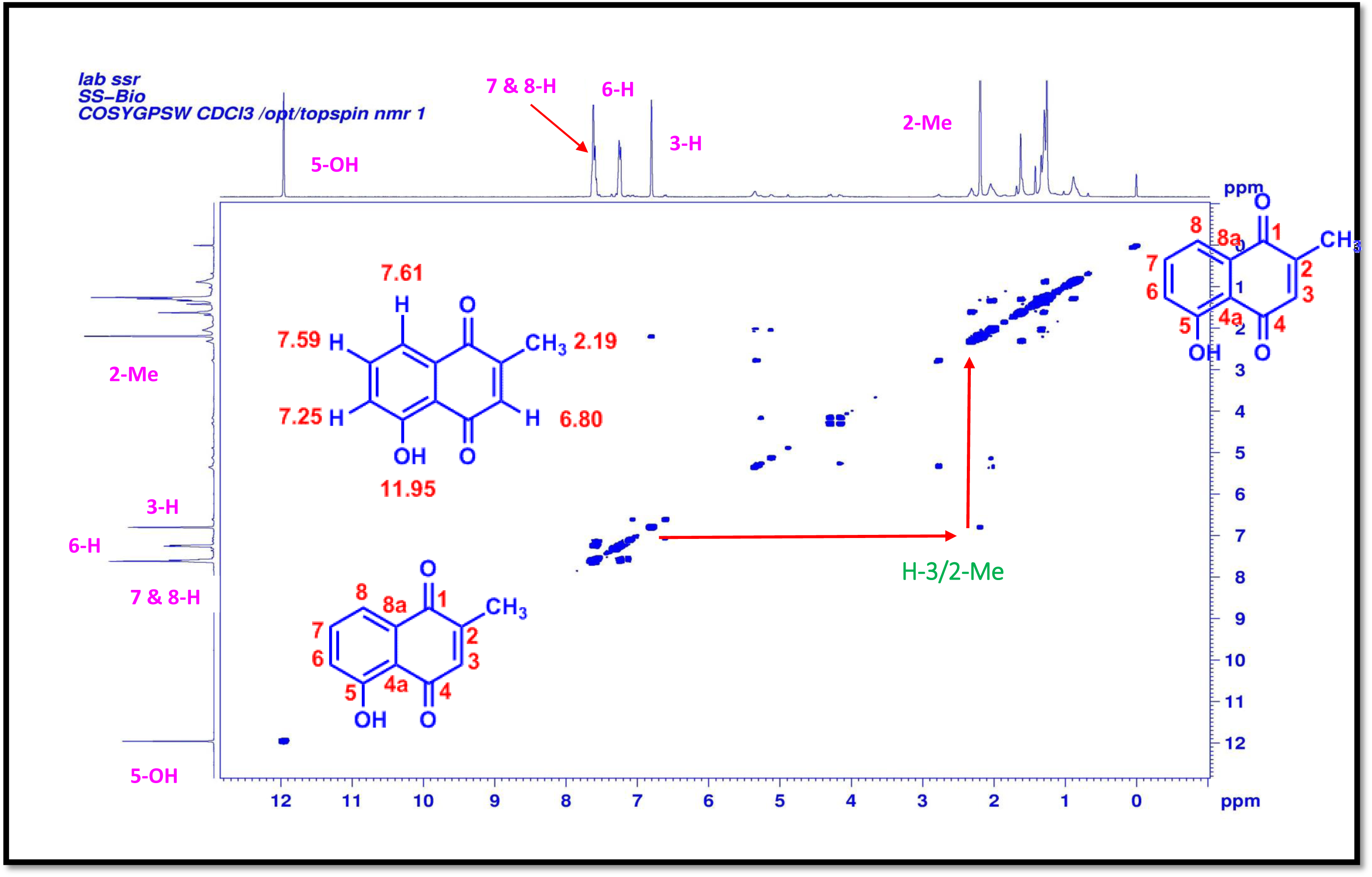
COSY (500 MHz, CDCl_3_) spectrum of plumbagin (1).

**Figure S7.**
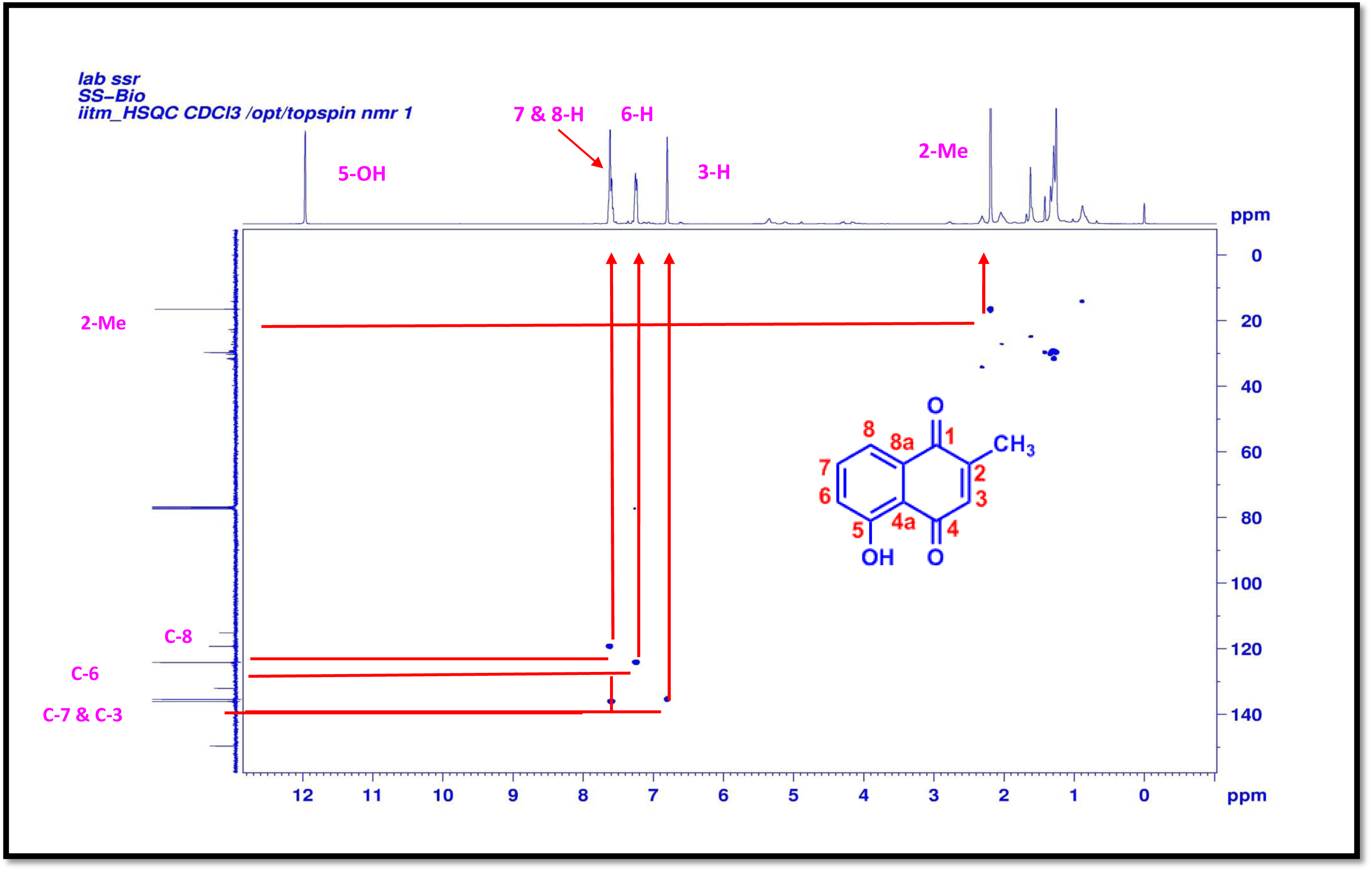
HSQC (500 MHz, CDCl_3_) spectrum of plumbagin (1).

**Figure S8.**
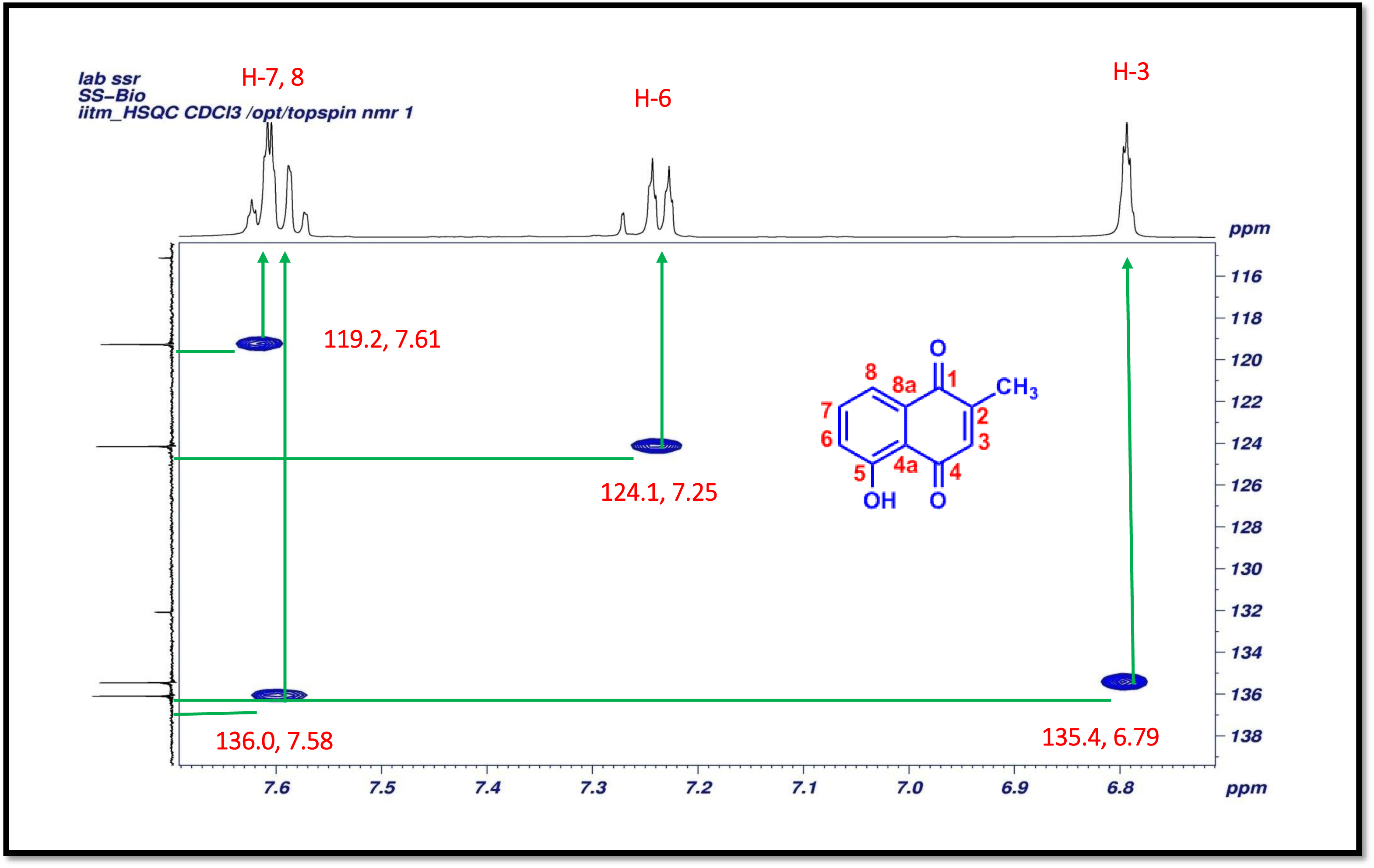
Expansion of HSQC (500 MHz, CDCl_3_) spectrum of plumbagin (1).

**Figure S9.**
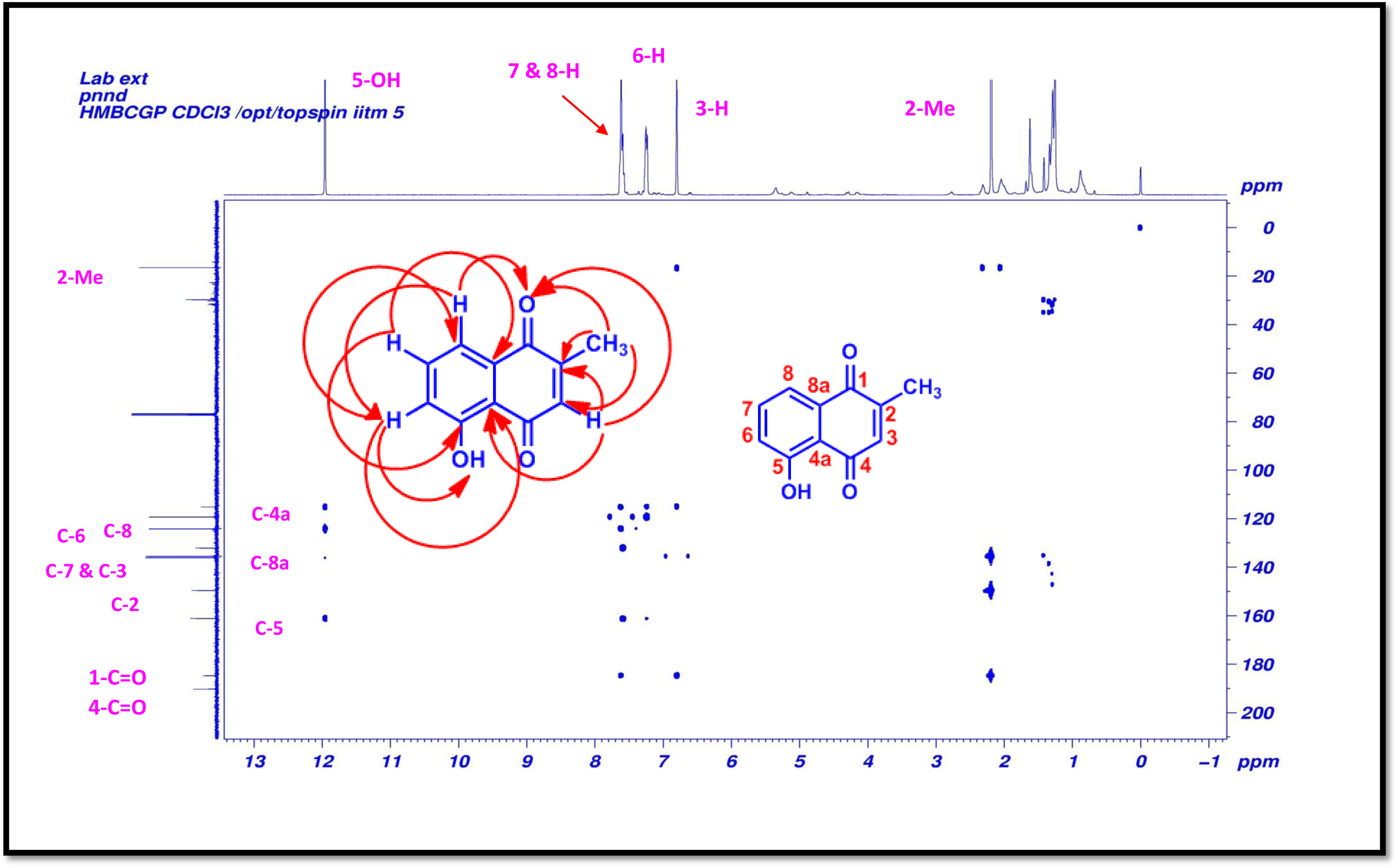
HMBC (500 MHz, CDCl_3_) spectrum of plumbagin (1).

**Figure S10.**
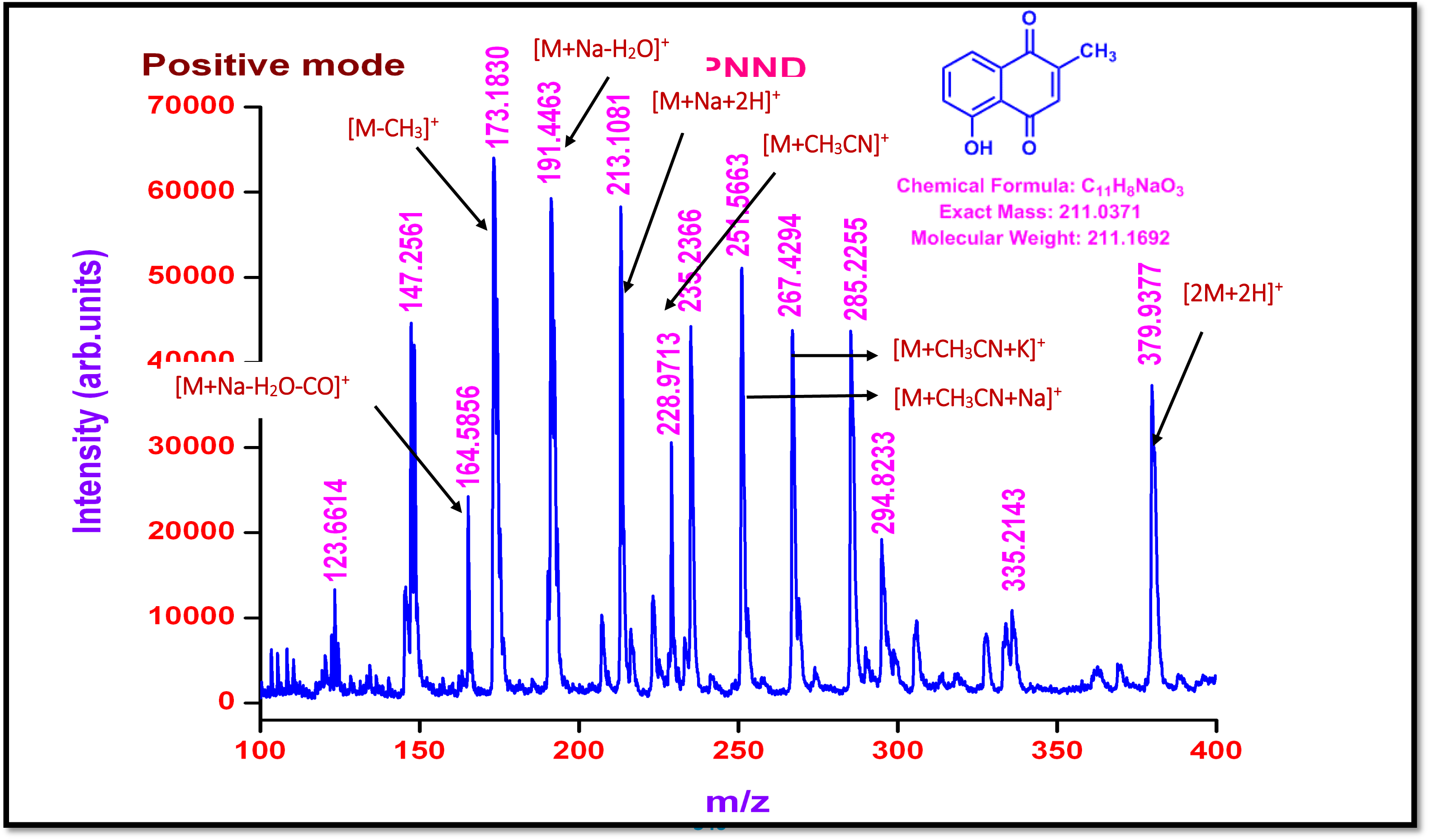
MALDI-TOF MS spectrum of plumbagin (1) (Positive mode).

**Figure S11.**
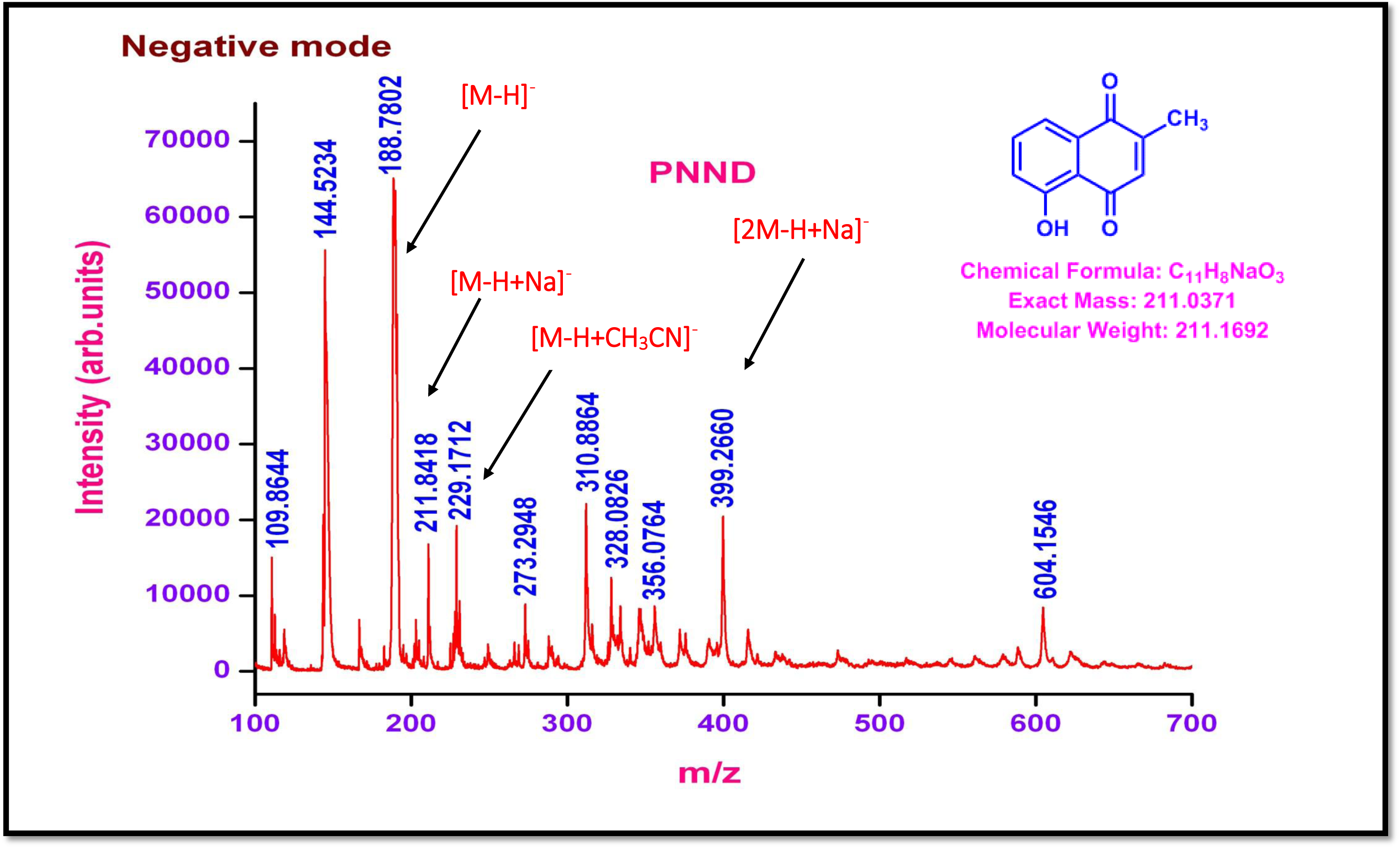
MALDI-TOF MS spectrum of plumbagin (1) (Negative mode).

**Figure S12.**
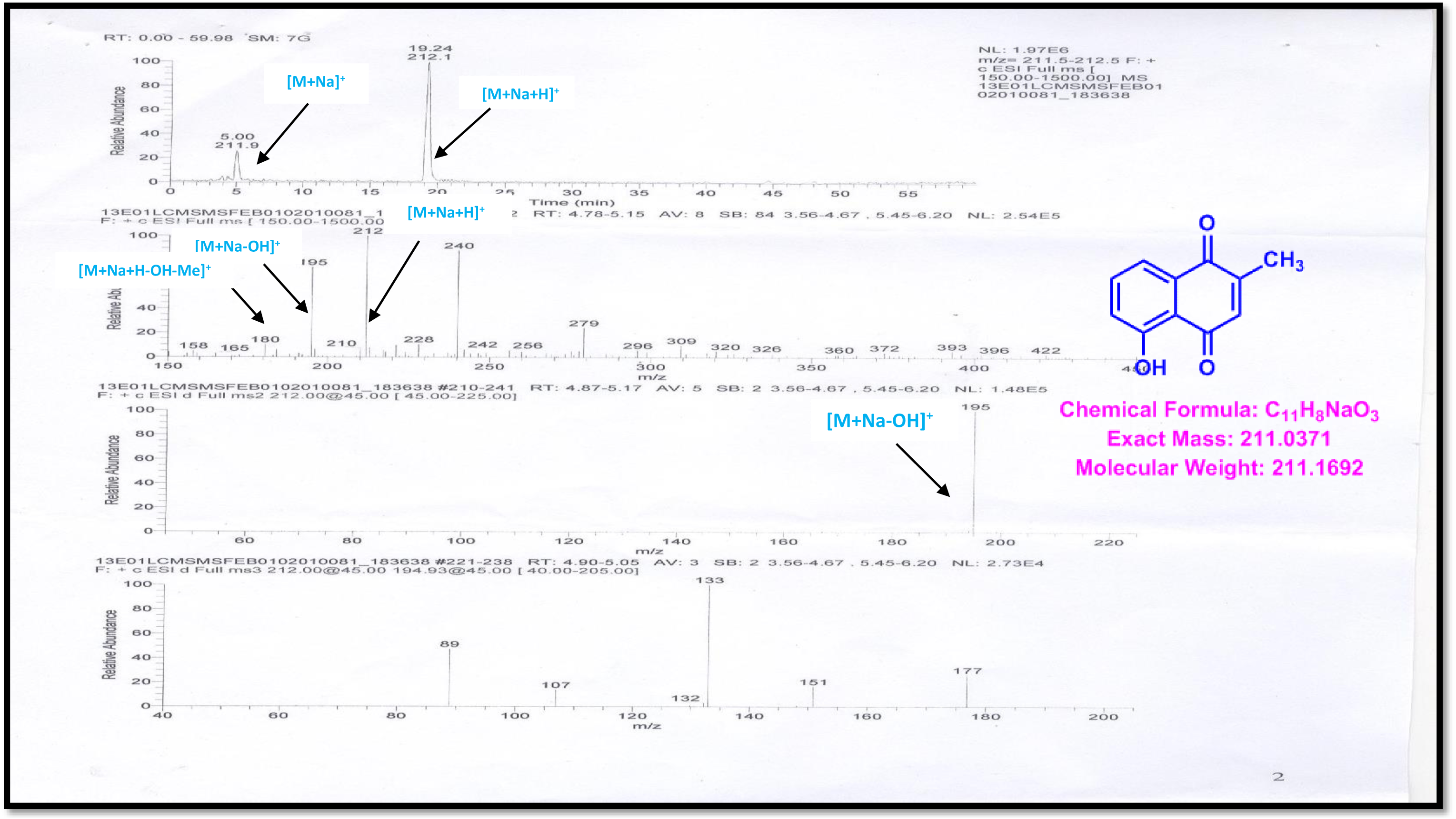
LC-EI-MS/MS spectrum of plumbagin (1).

**Figure S13.**
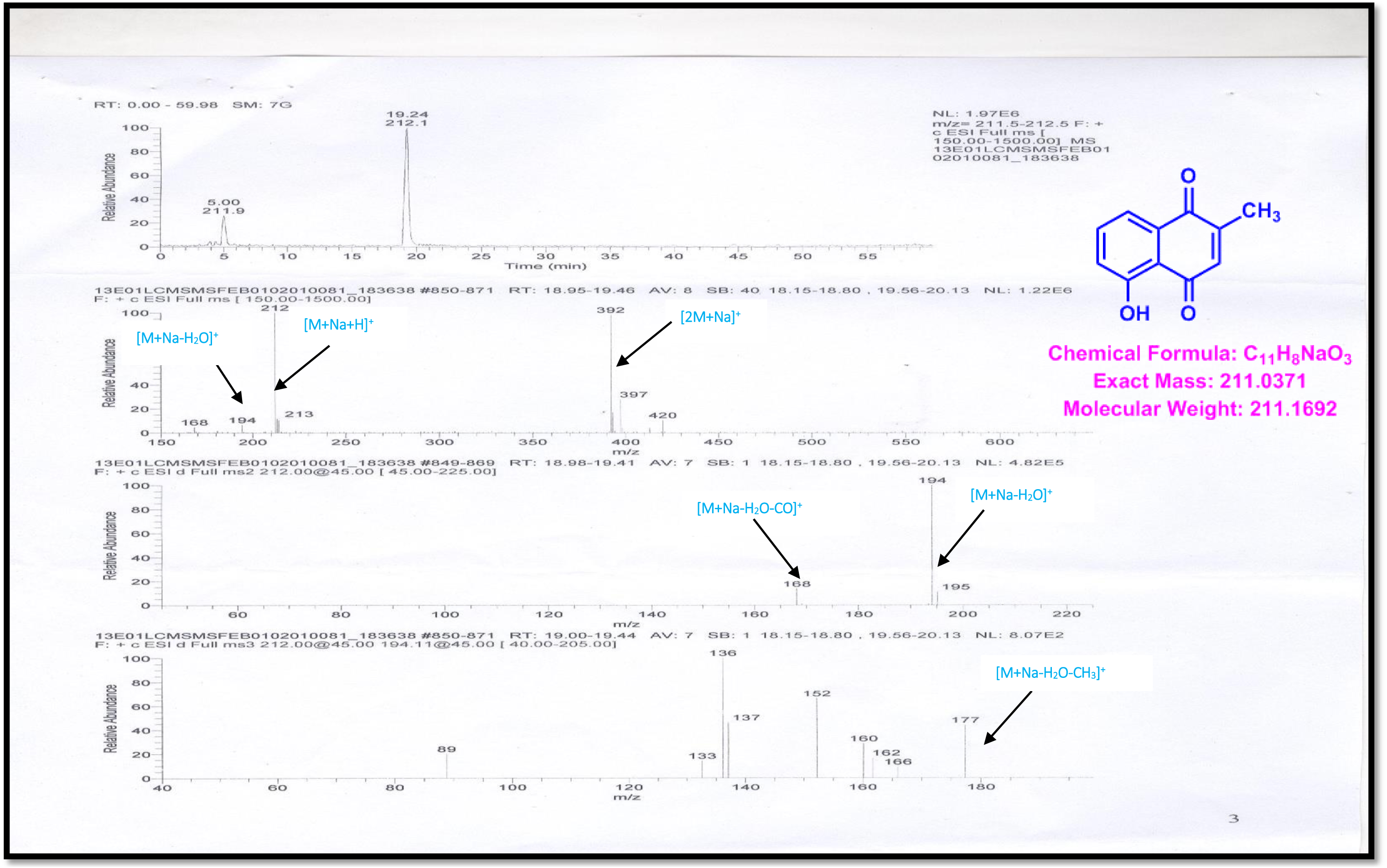
LC-ESI-MS/MS spectrum of plumbagin (1).

**Figure S14.**
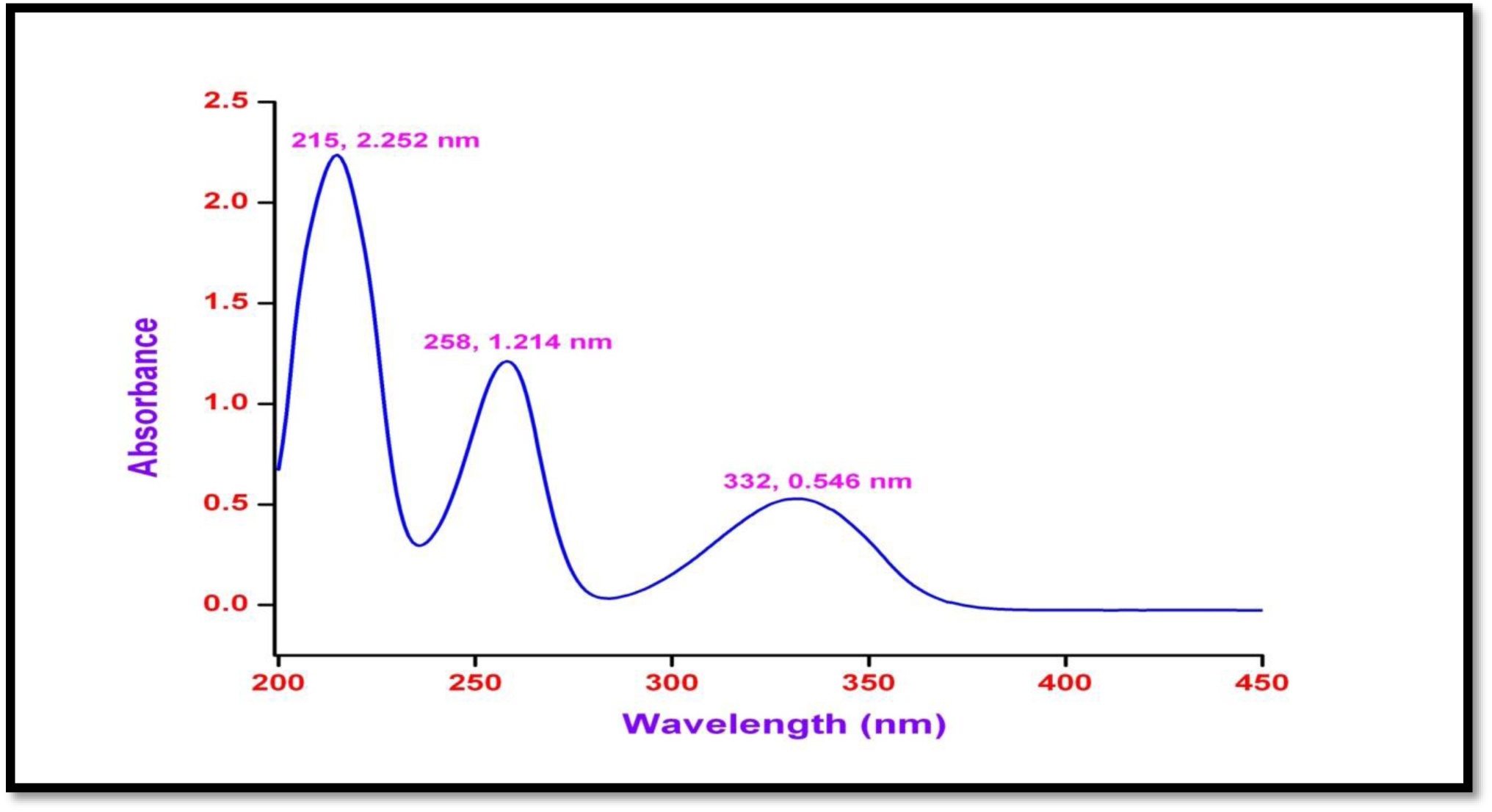
UV/Vis spectrum of cis-isoshinonolone (2) in methanol. UV: (MeOH) λ max (log ε) 216 (2.252) 258 (1.214), 332 (0.546) nm

**Figure S15.**
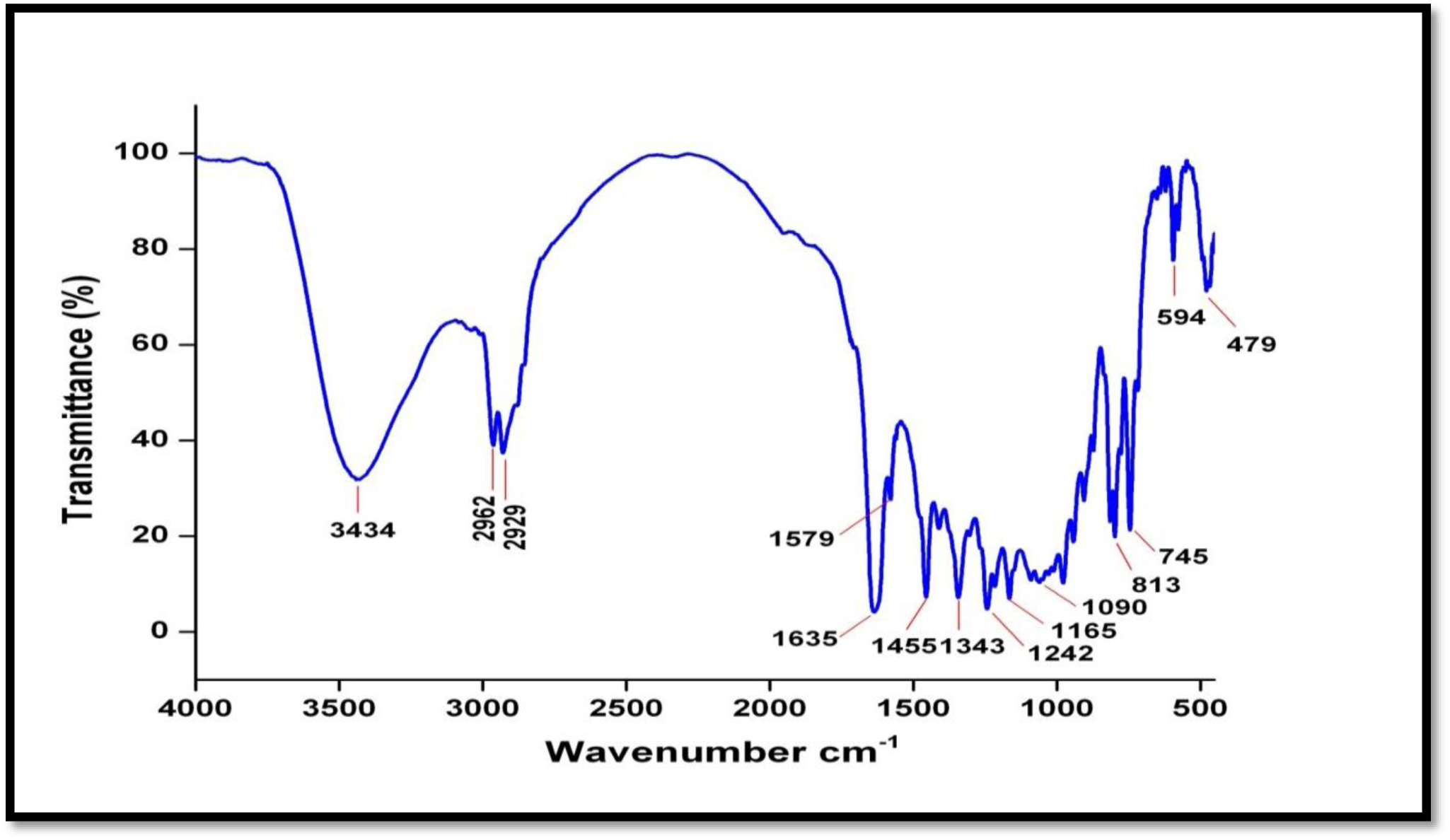
IR spectrum of cis-isoshinonolone (2).

**Figure S16.**
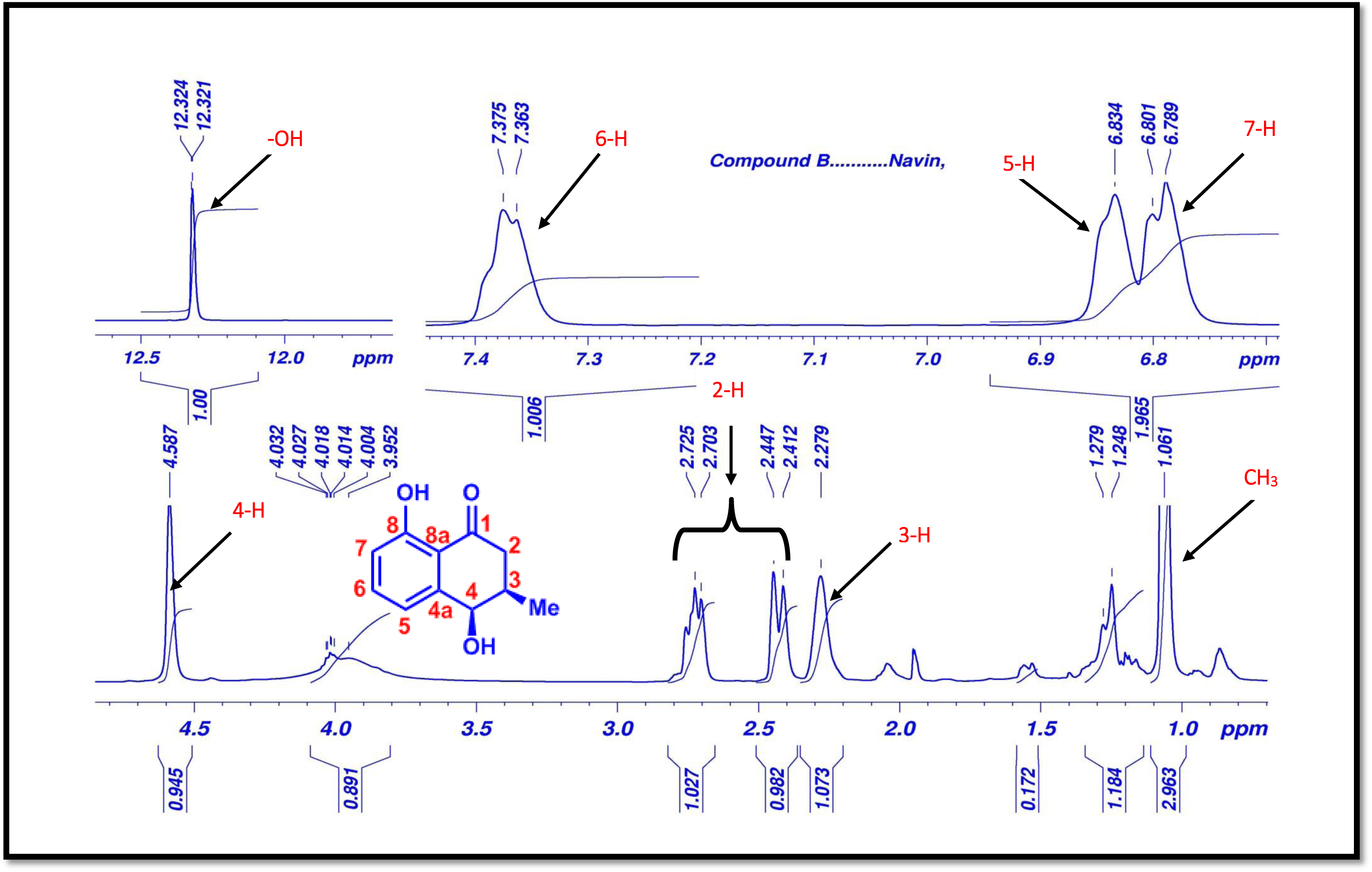
^1^H-NMR (500 MHz, CDCl_3_) spectrum of cis-isoshinonolone (2).

**Figure S17.**
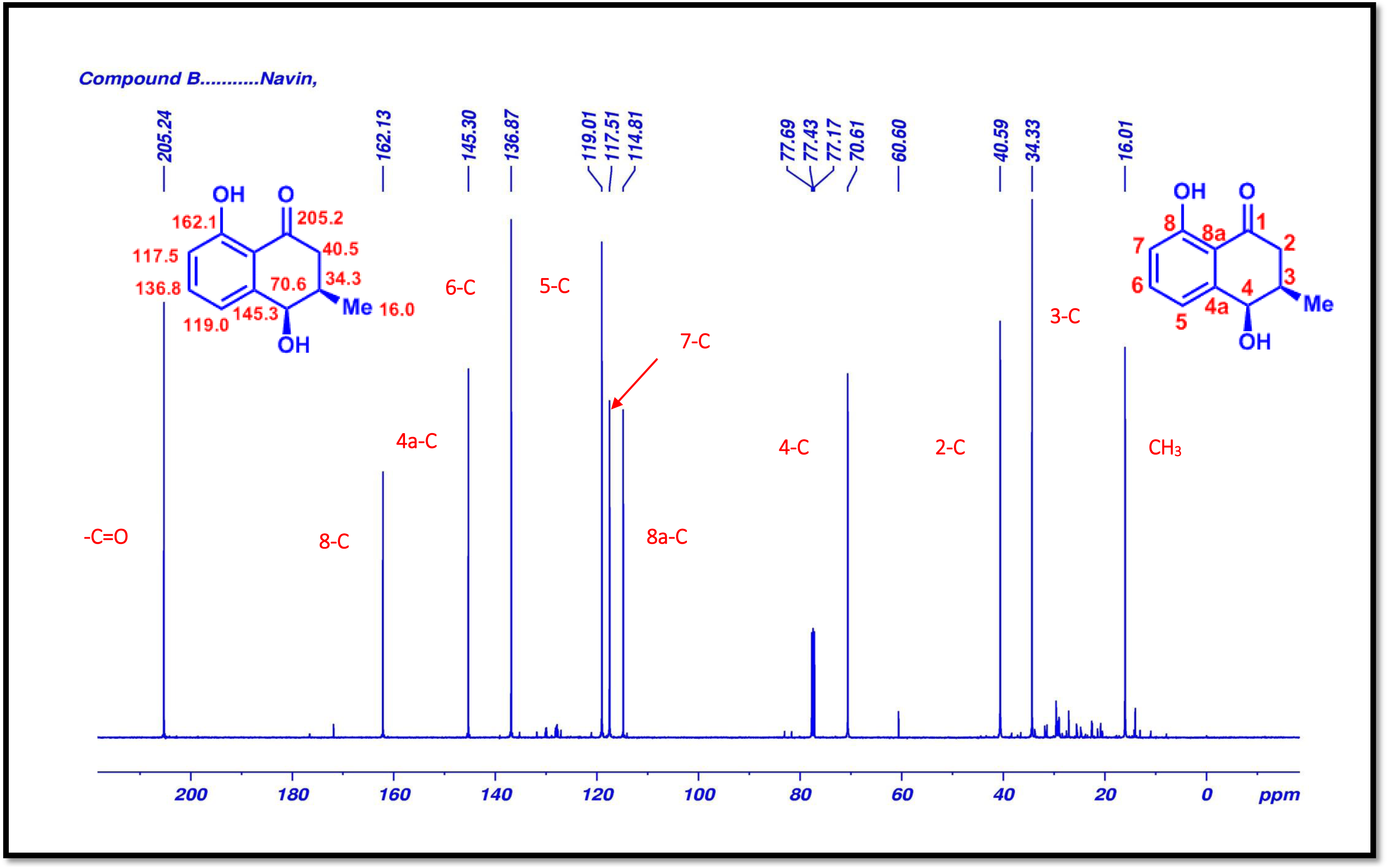
^13^C-NMR (125 MHz, CDCl_3_) spectrum of cis-isoshinonolone (2).

**Figure S18.**
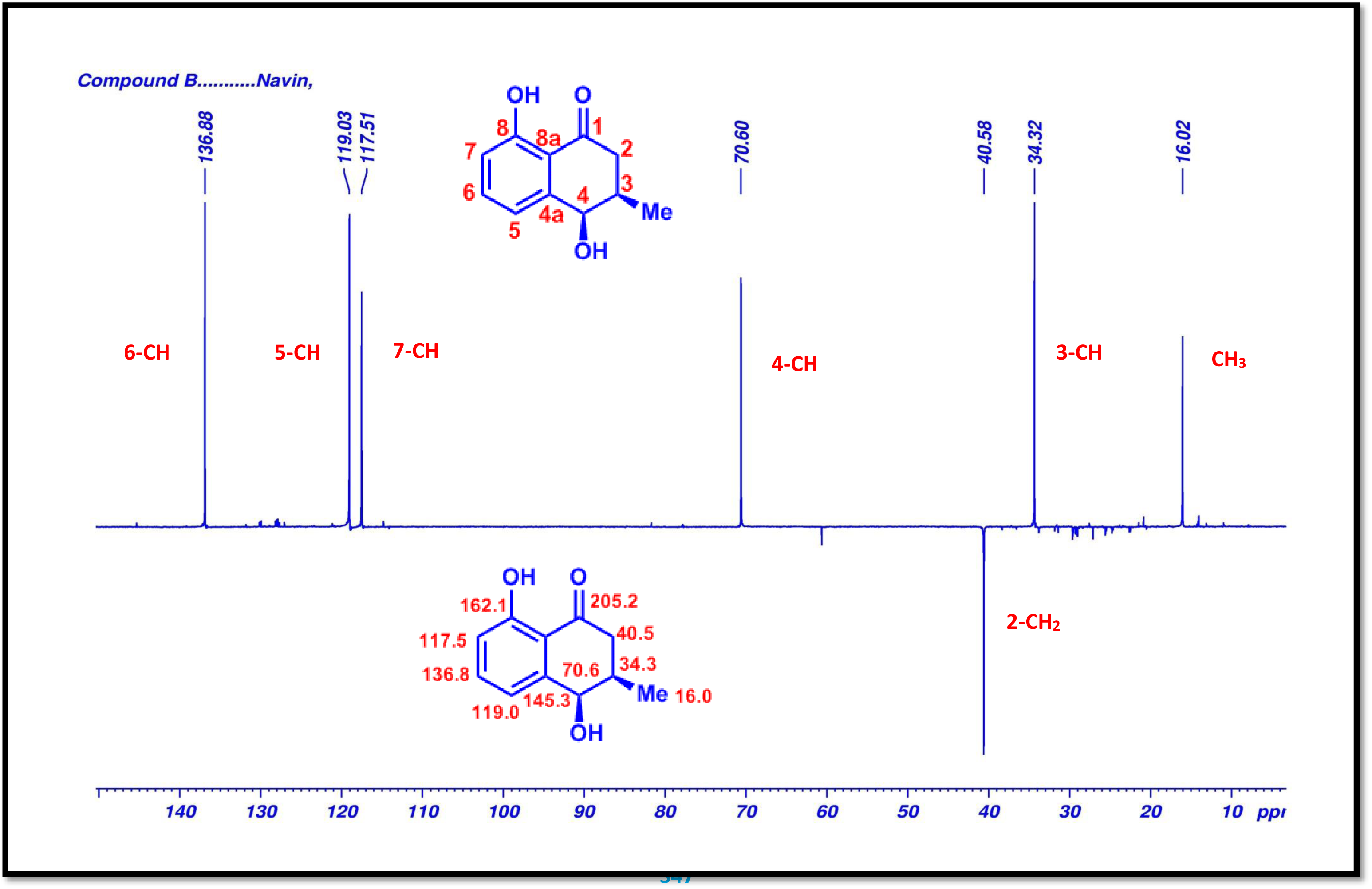
DEPT135 (500 MHz, CDCl_3_) Spectrum of cis-isoshinonolone (2).

**Figure S19.**
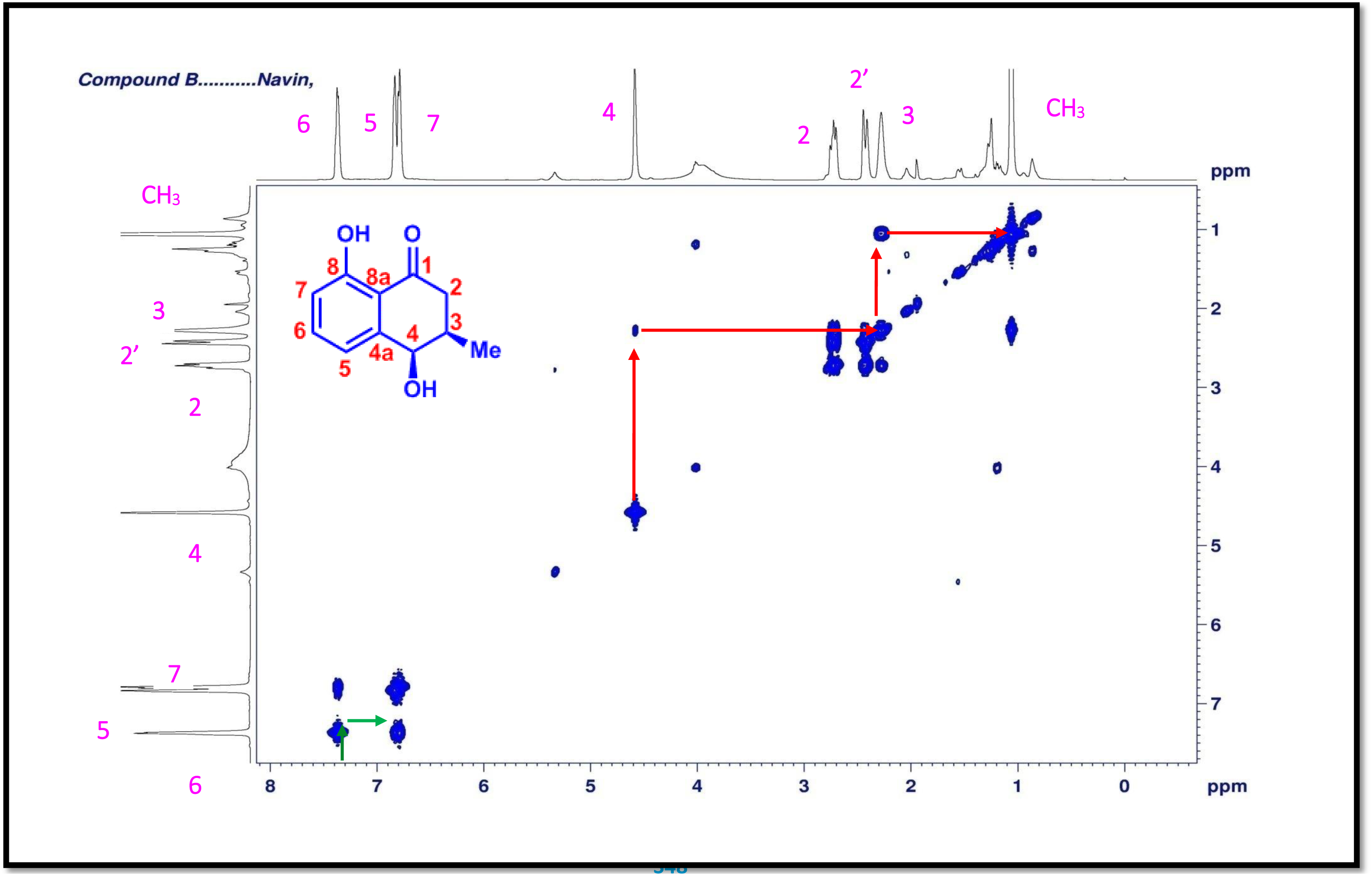
COSY (500 MHz, CDCl_3_) spectrum of cis-isoshinonolone (2).

**Figure S20.**
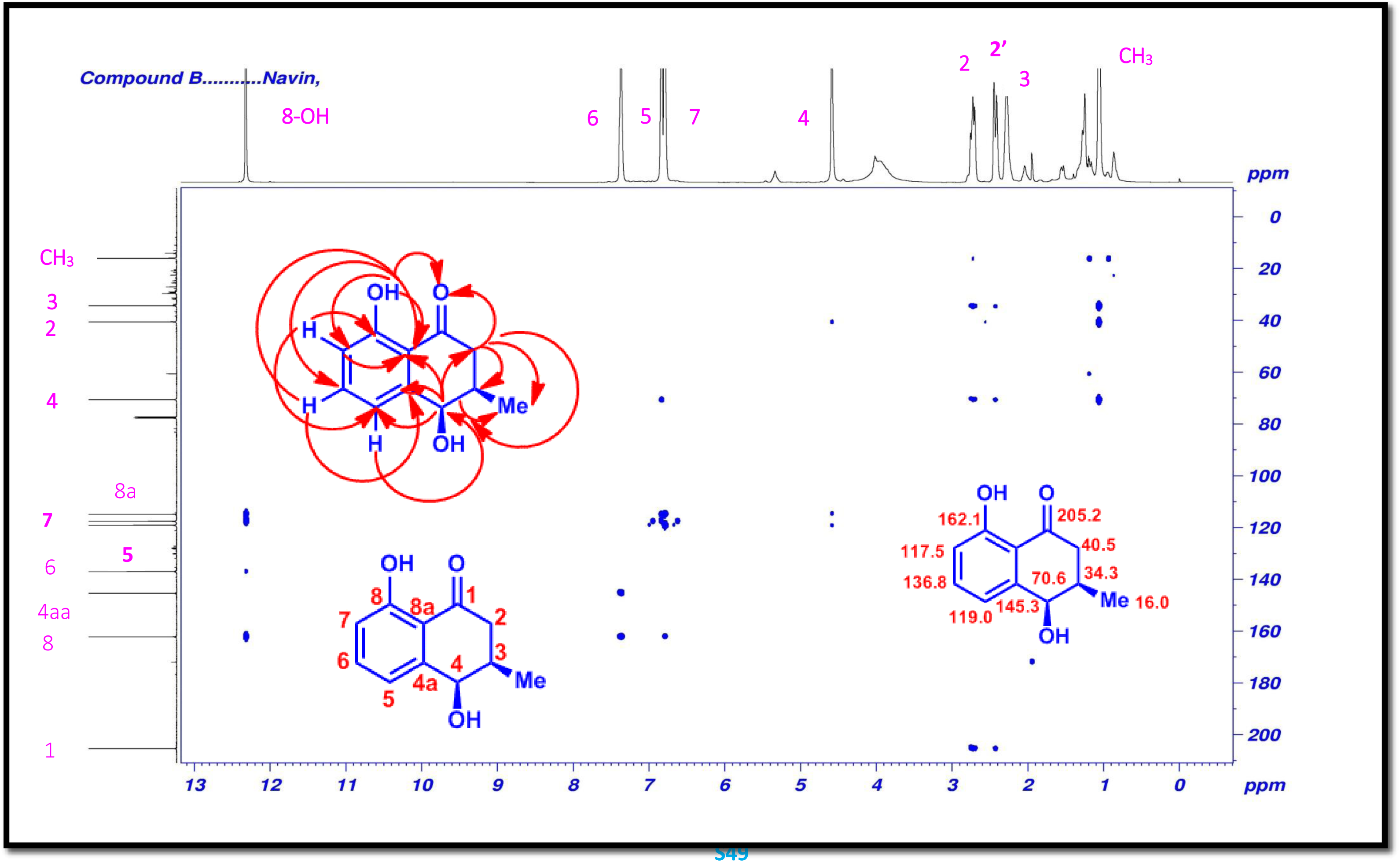
HMBC (500 MHz, CDCl_3_) spectrum of cis-isoshinonolone (2).

**Figure S21.**
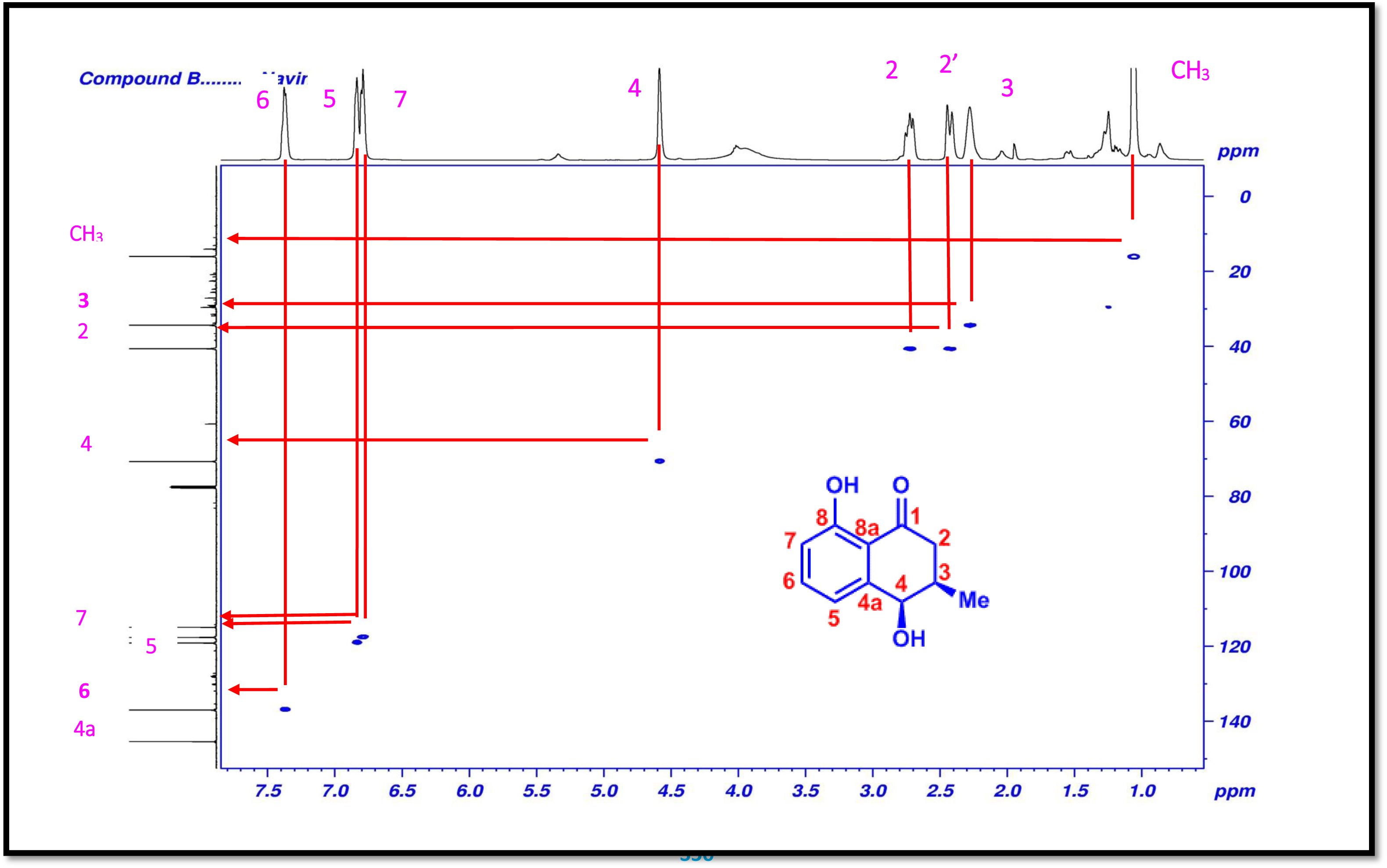
HSQC (500 MHz, CDCl_3_) spectrum of cis-isoshinonolone (2).

**Figure S22.**
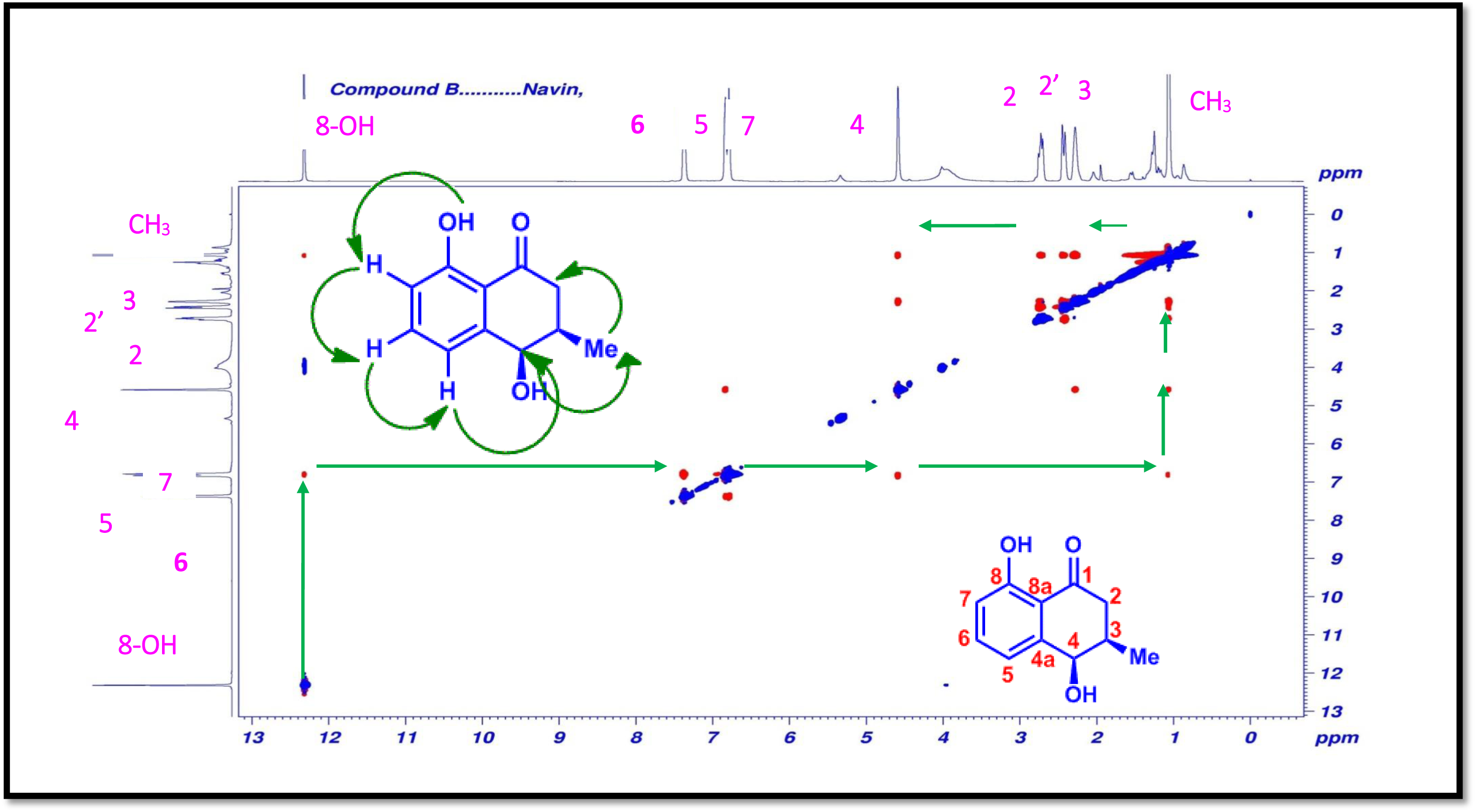
NOSEY (500 MHz,) spectrum of cis-isoshinonolone (2).

**Figure S23.**
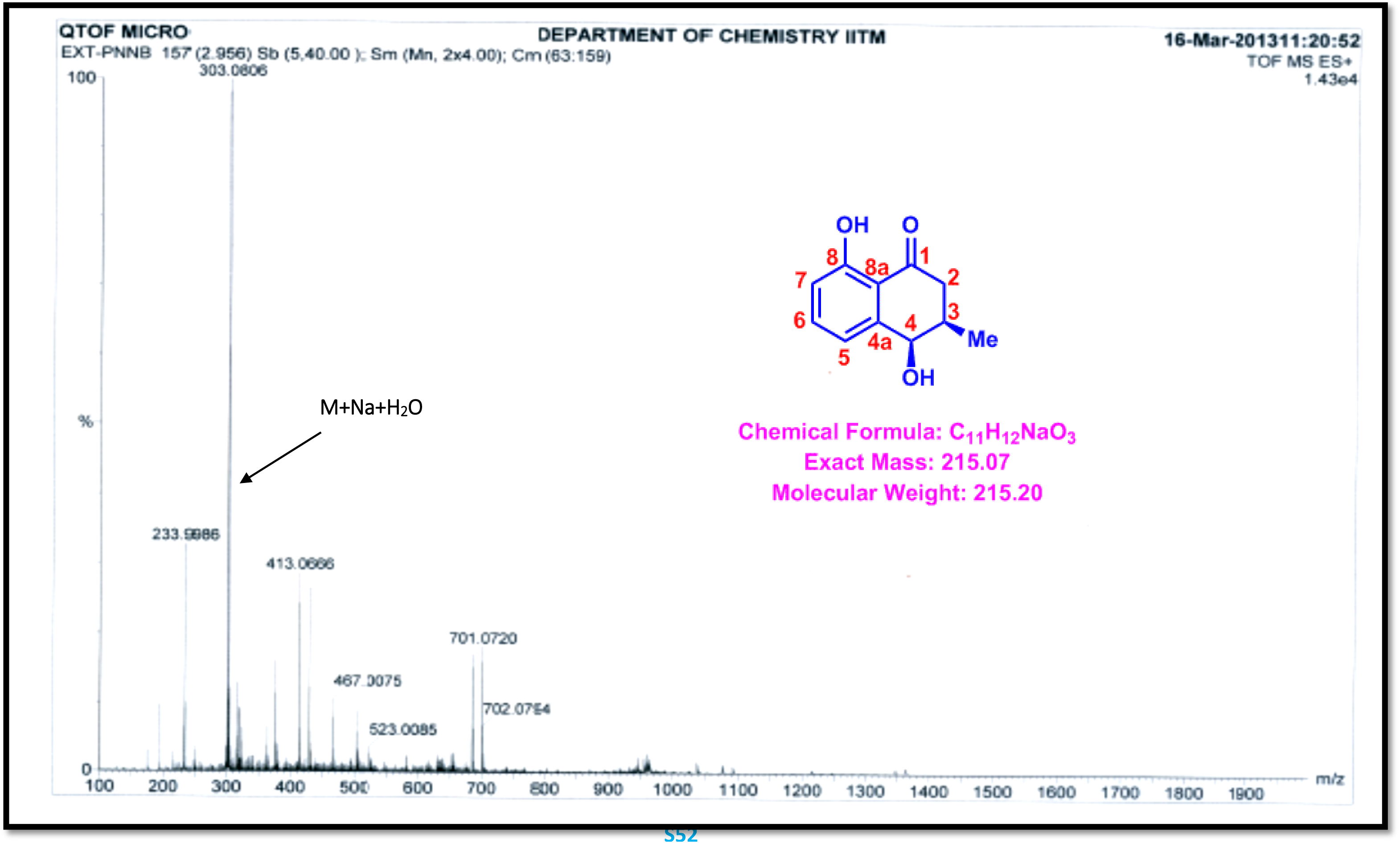
ESI-QTOF-MS spectrum of cis-isoshinonolone (2).

**Figure S24.**
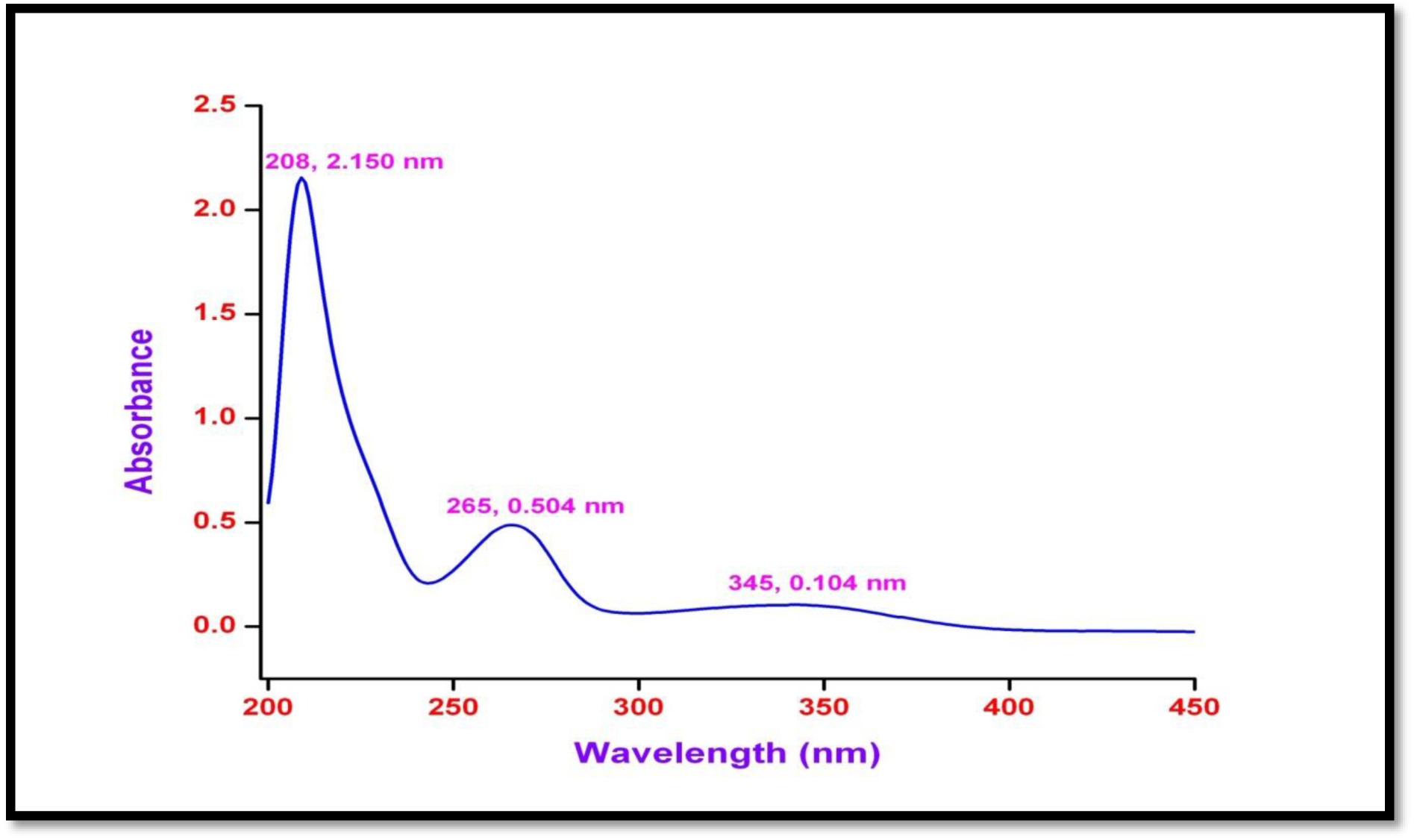
UV/Vis spectrum of 3’-o-β-glucopyranosyl plumbagic acid (3) in methanol. UV: (MeOH) λ *_max_* (log ε) 208 (2.150) 265 (0.504), 354 (0.104) nm

**Figure S25.**
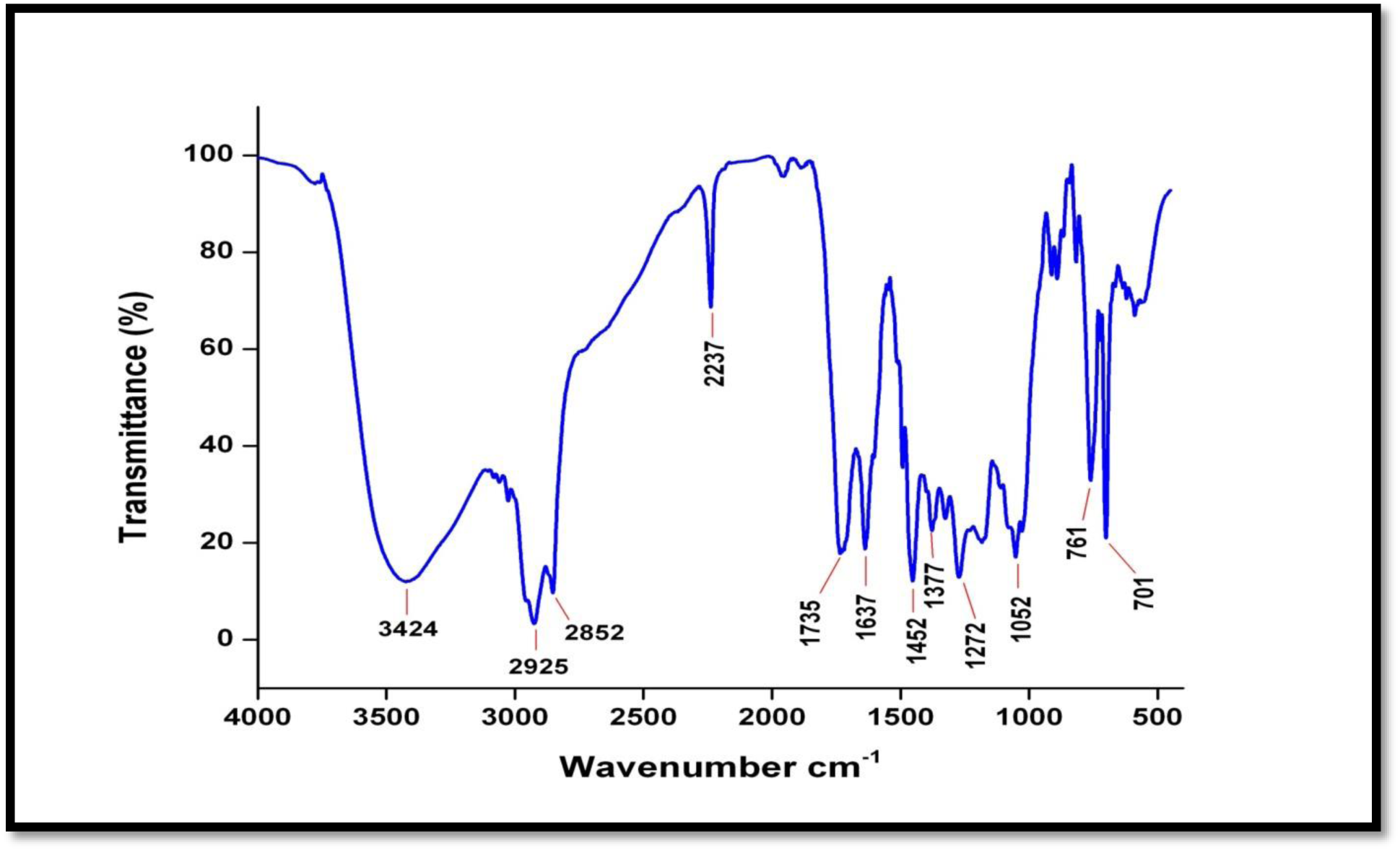
IR spectrum of 3’-o-β-glucopyranosyl plumbagic acid (3).

**Figure S26.**
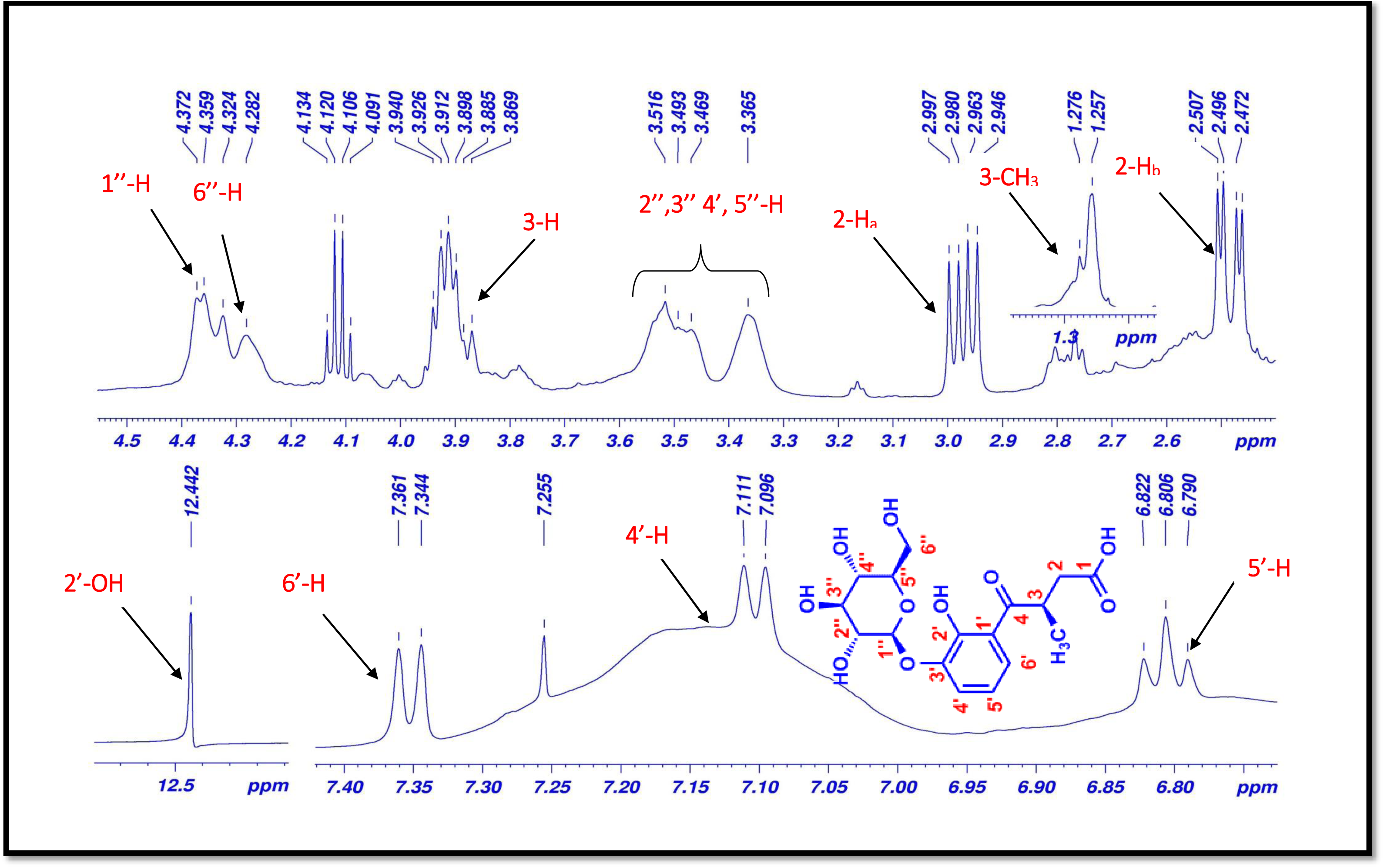
^1^H-NMR (500 MHz, CDCl_3_) spectrum of 3’-o-β-glucopyranosyl plumbagic acid (3).

**Figure S27.**
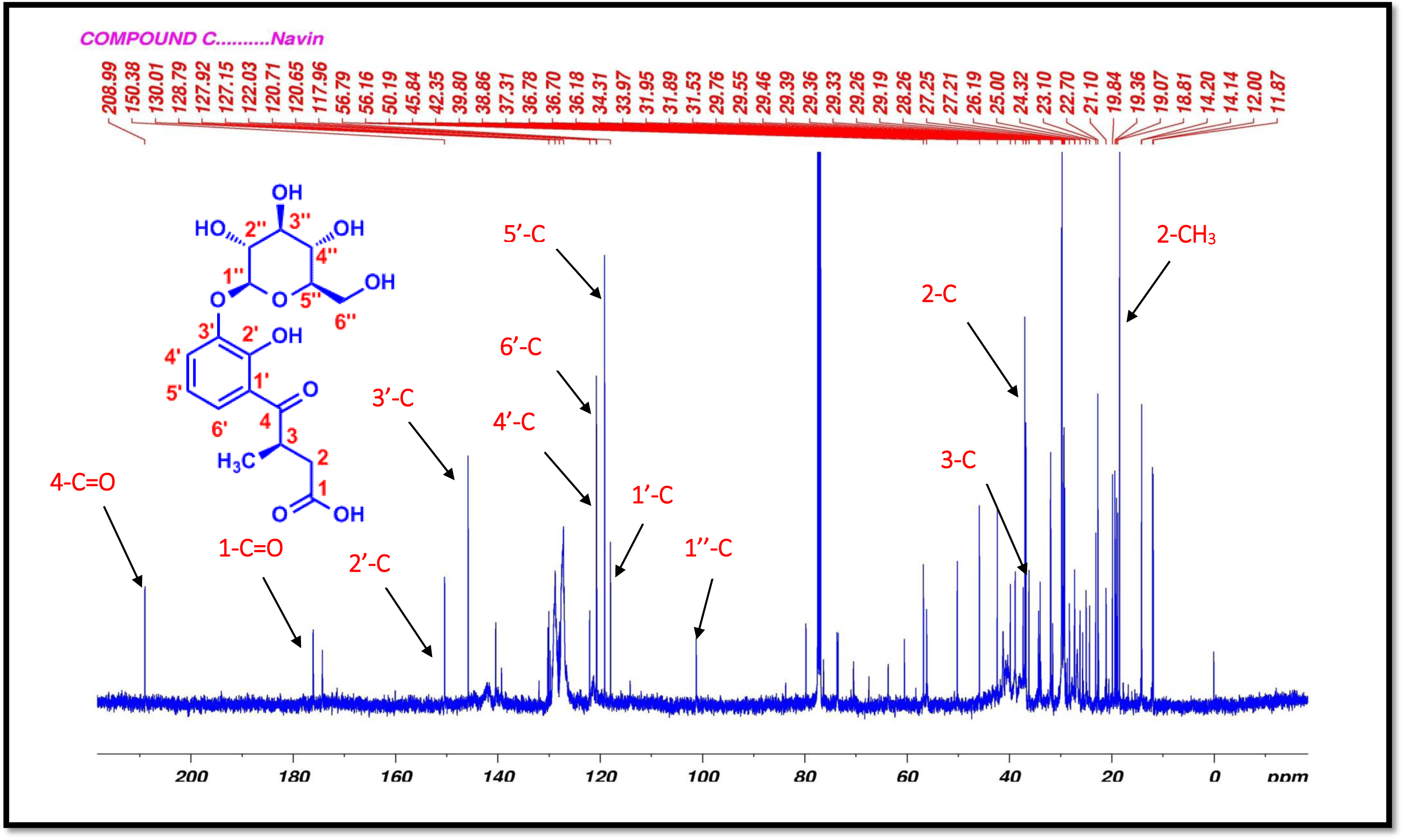
^13^C-NMR (125 MHz, CDCl_3_) spectrum of 3’-o-β-glucopyranosyl plumbagic acid (3).

**Figure S28.**
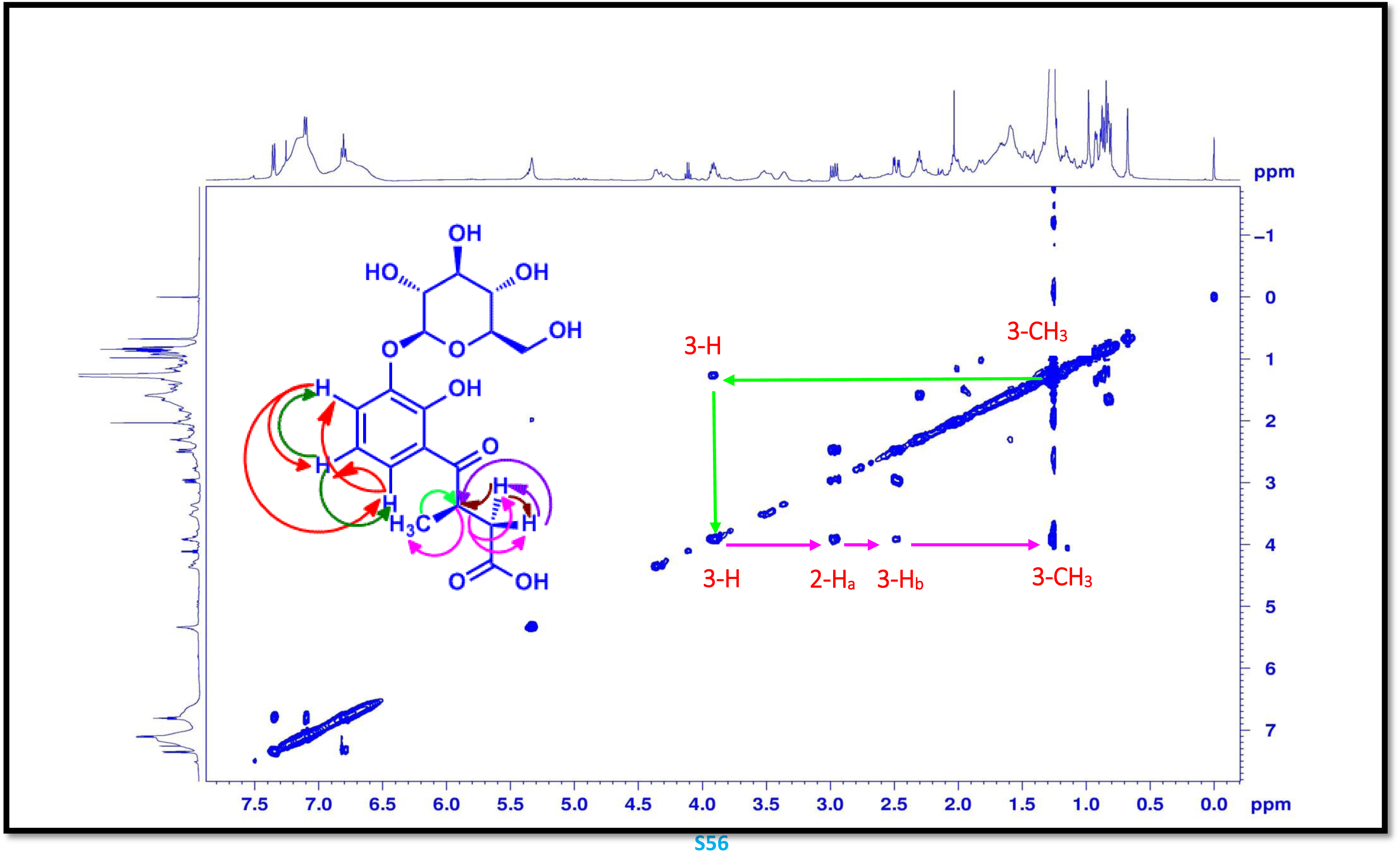
COSY (125 MHz, CDCl_3_) spectrum of 3’-o-β-glucopyranosyl plumbagic acid (3).

**Figure S29.**
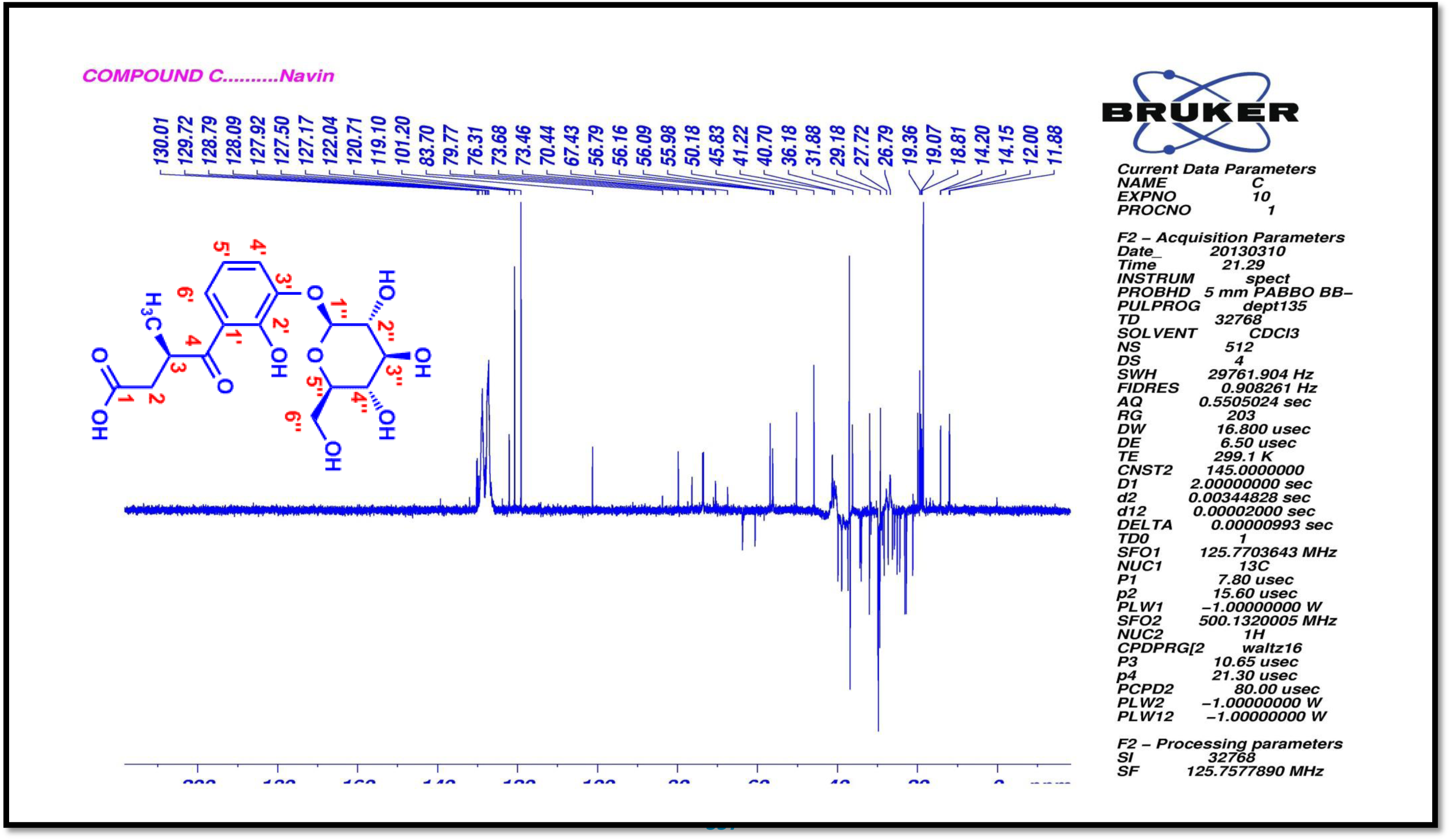
DEPT135 (500 MHz, CDCl_3_) spectrum of 3’-o-β-glucopyranosyl plumbagic acid (3).

**Figure S30.**
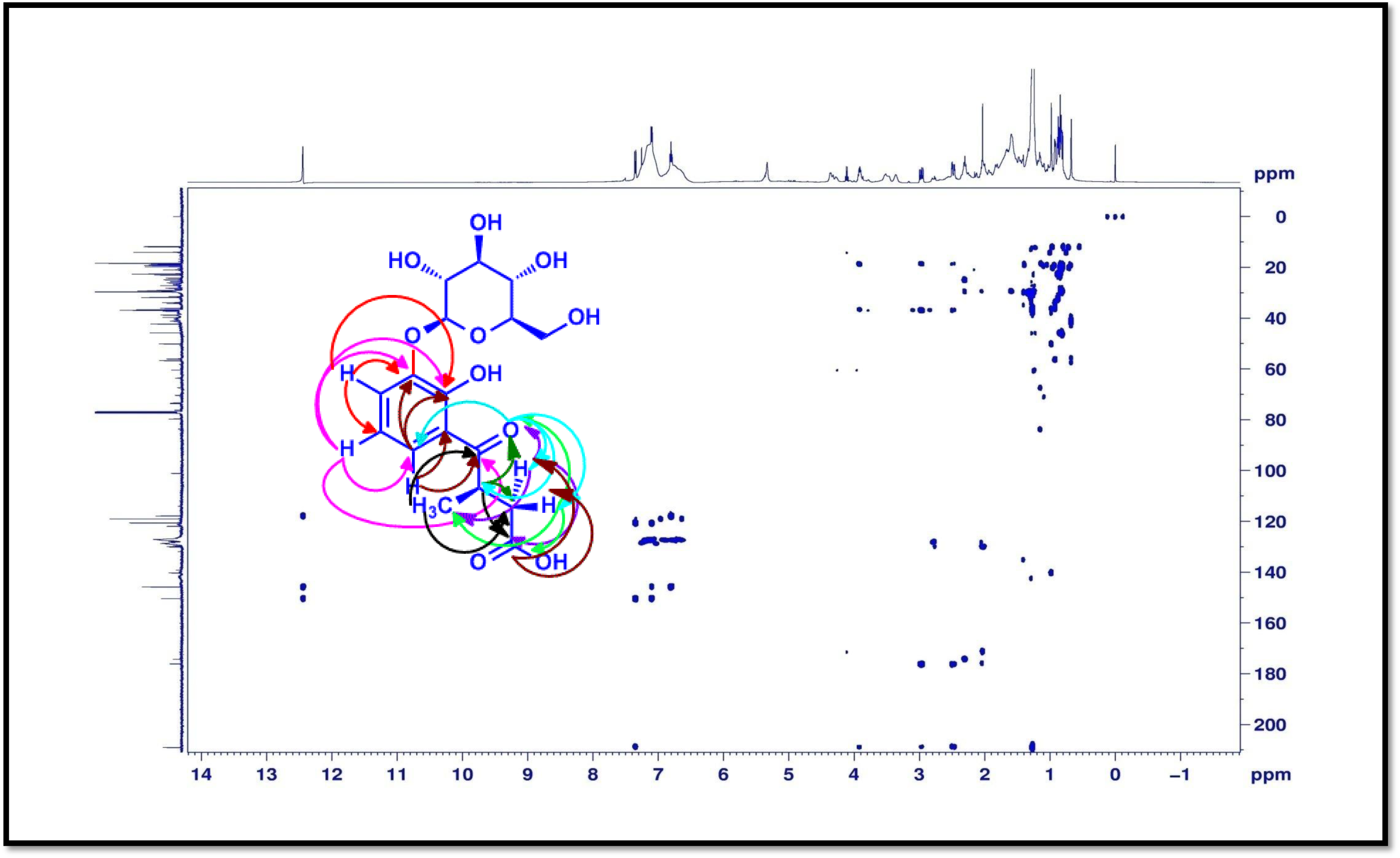
HMBC (500 MHz, CDCl_3_) spectrum of 3’-o-β-glucopyranosyl plumbagic acid (3).

**Figure S31.**
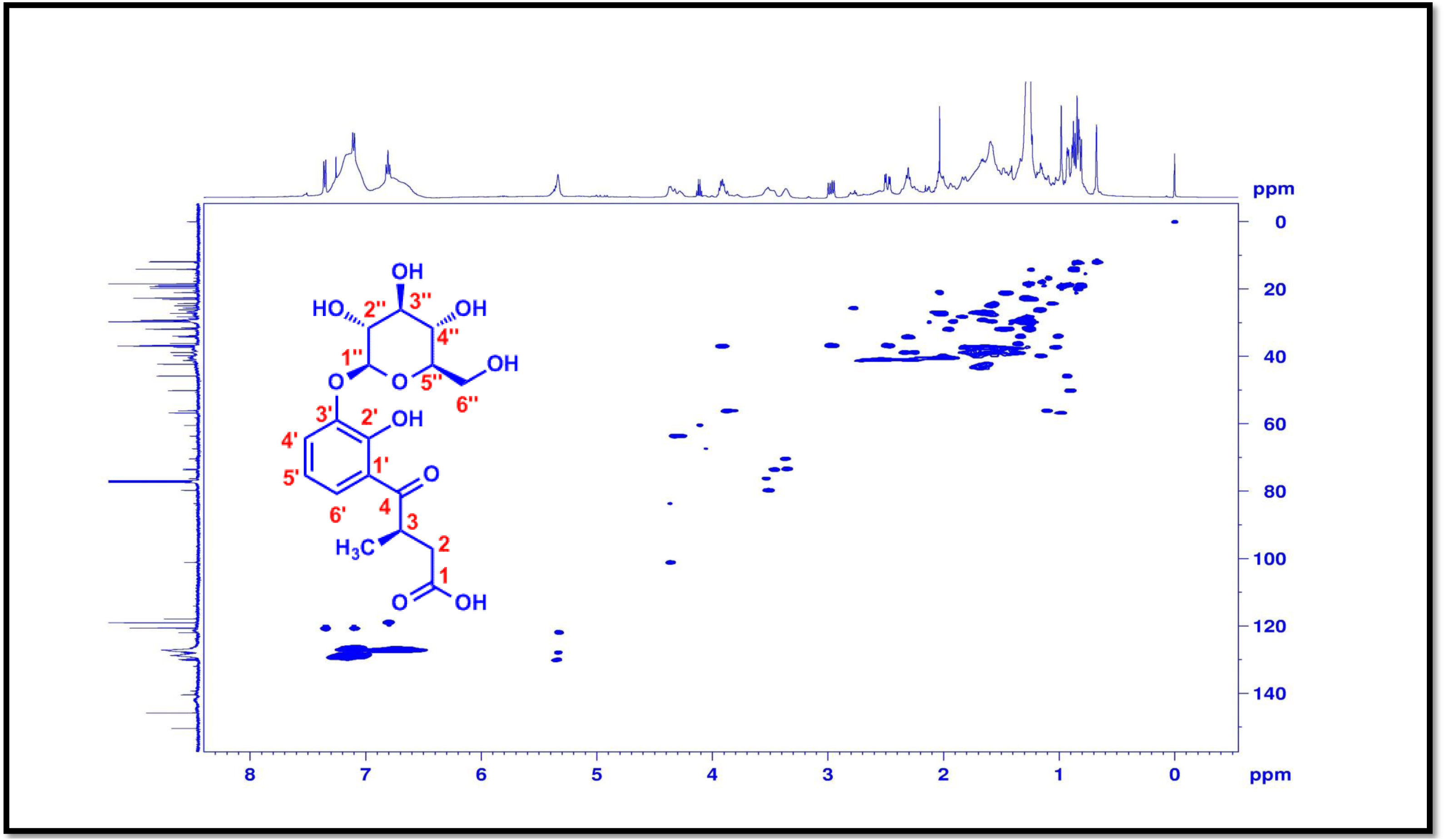
HSQC (500 MHz, CDCl_3_) spectrum of 3’-o-β-glucopyranosyl plumbagic acid (3).

**Figure S32.**
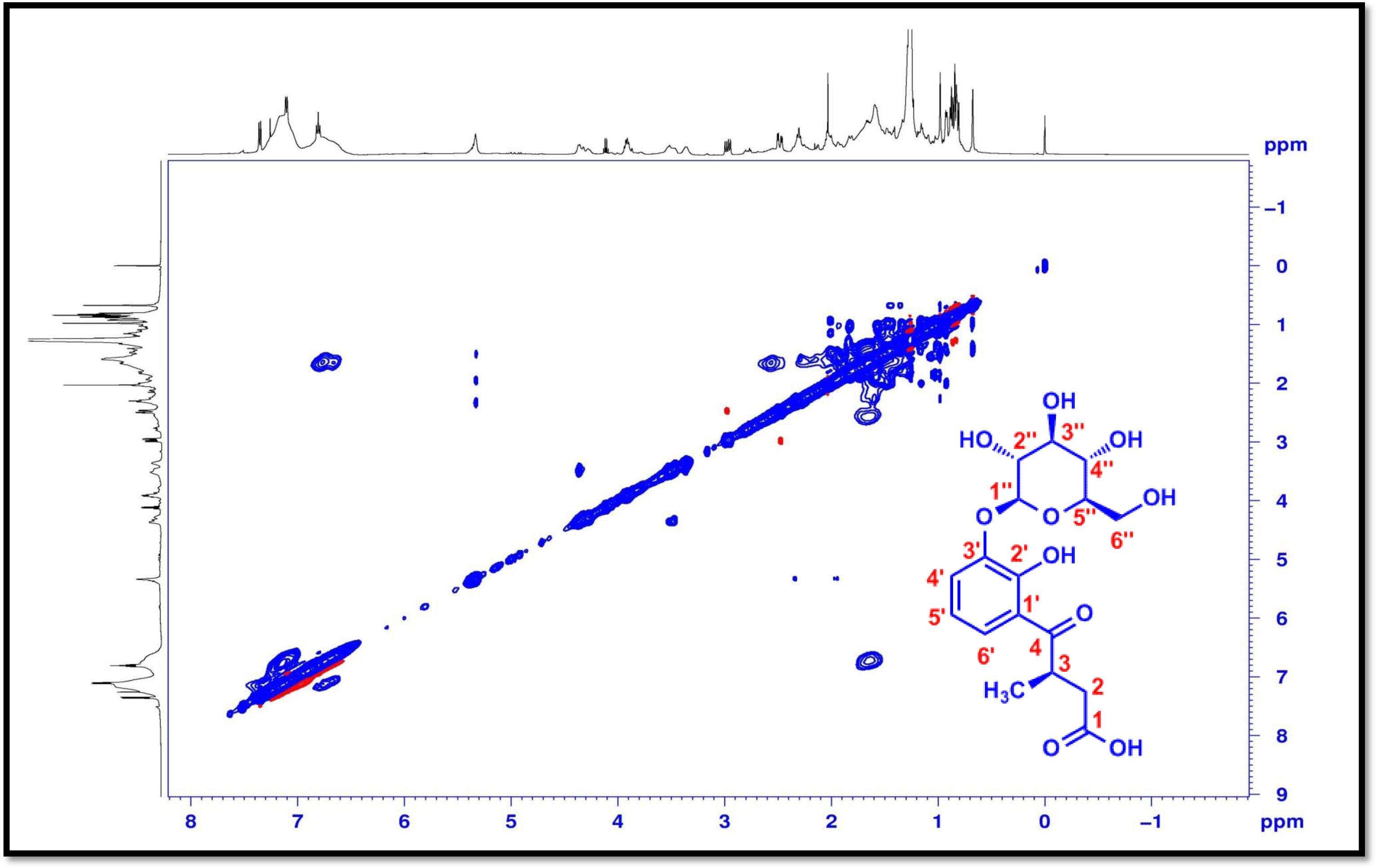
NOSEY (500 MHz, CDCl_3_) spectrum of 3’-o-β-glucopyranosyl plumbagic acid (3).

**Figure S33.**
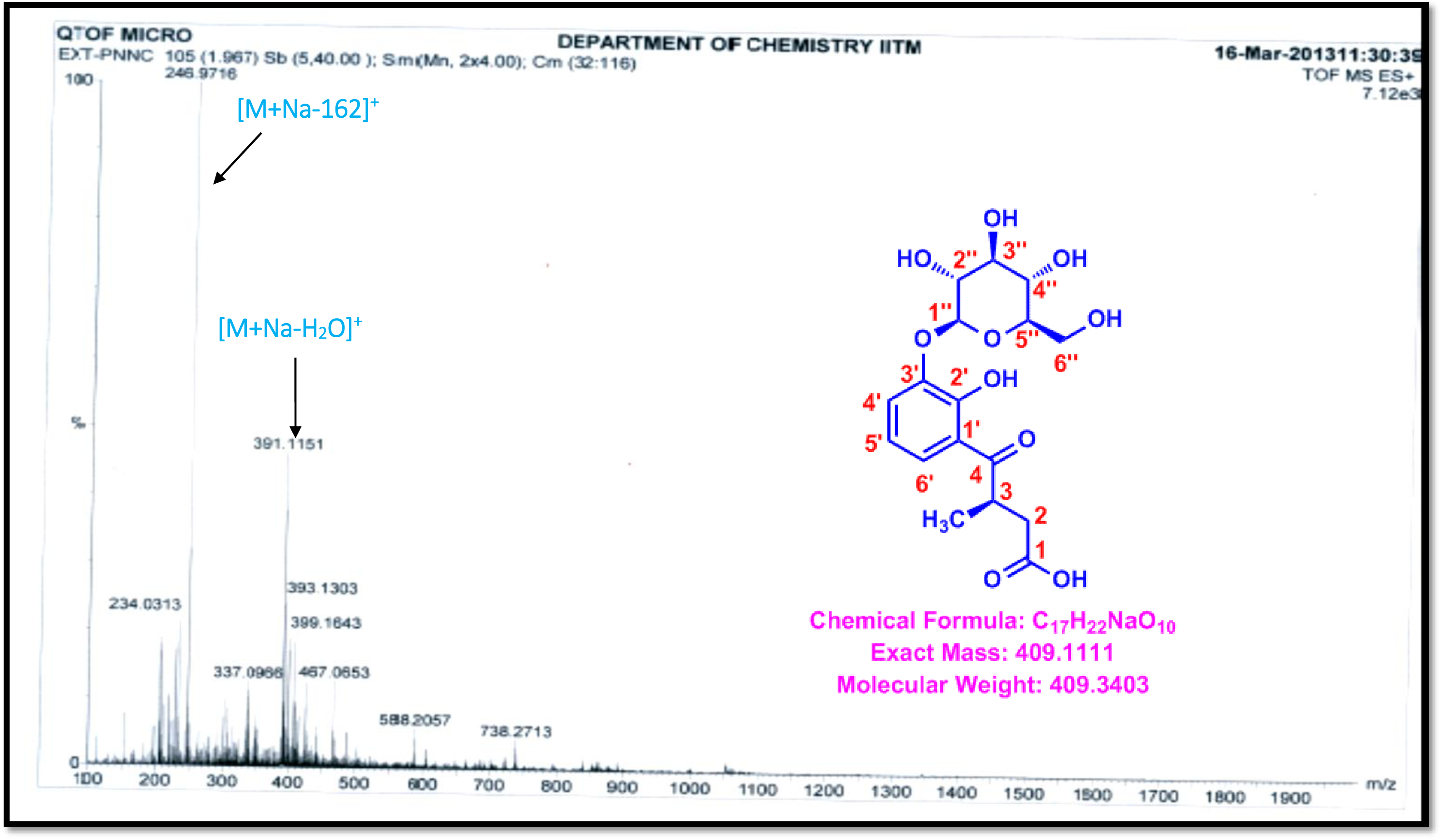
ESI-MS spectrum of 3’-o-β-glucopyranosyl plumbagic acid (3).

